# Crystal structure of *Arabidopsis thaliana* HPPK/DHPS, a bifunctional enzyme and target of the herbicide asulam

**DOI:** 10.1101/2021.11.10.468163

**Authors:** Grishma Vadlamani, Kirill V. Sukhoverkov, Joel Haywood, Karen J. Breese, Mark F. Fisher, Keith A. Stubbs, Charles S. Bond, Joshua S. Mylne

## Abstract

Herbicides are vital for modern agriculture, but their utility is threatened by genetic or metabolic resistance in weeds as well as heightened regulatory scrutiny. Of the known herbicide modes of action, 6-hydroxymethyl-7,8-dihydropterin synthase (DHPS) which is involved in folate biosynthesis, is targeted by just one commercial herbicide, asulam. A mimic of the substrate *para*-aminobenzoic acid, asulam is chemically similar to sulfonamide antibiotics – and while still in widespread use, asulam has faced regulatory scrutiny. With an entire mode of action represented by just one commercial agrochemical, we sought to improve the understanding of its plant target. Here we solve a 2.6 Å resolution crystal structure for *Arabidopsis thaliana* DHPS that is conjoined to 6-hydroxymethyl-7,8-dihydropterin pyrophosphokinase (HPPK) and reveal a strong structural conservation with bacterial counterparts at the sulfonamide-binding pocket of DHPS. We demonstrate asulam and the antibiotics sulfacetamide and sulfamethoxazole have herbicidal as well as antibacterial activity and explore the structural basis of their potency by modelling these compounds in mitochondrial HPPK/DHPS. Our findings suggest limited opportunity for the rational design of plant selectivity from asulam and that pharmacokinetic or delivery differences between plants and microbes might be the best approaches to safeguard this mode of action.

## Introduction

Asulam is the only commercial herbicide that targets the folate biosynthesis pathway. Although humans obtain folate from their diet, plants and microorganisms synthesise tetrahydrofolic acid (THF) using 6-hydroxymethylpterin, glutamic acid and *para*-aminobenzoic acid (*p*-ABA) as precursors (Hossain et al., 2004). THF and its derivatives: methyl-THF, methenyl-THF and formyl-THF are involved in one-carbon transfer reactions and are part of the methylation cycle and the biosynthesis of DNA and amino acids (Hanson and Gregory, 2002; Hanson and Gregory, 2011). In plants THF is synthesised by the sequential activities of five mitochondrial enzymes, namely 6-hydroxymethyl-7,8-dihydropterin pyrophosphokinase (HPPK), 6-hydroxymethyl-7,8-dihydropterin synthase (DHPS), dihydrofolate synthetase (DHFS), dihydrofolate reductase (DHFR), and folylpolyglutamate synthetase (FPGS) (Hanson and Gregory, 2011). HPPK initiates tetrahydrofolate biosynthesis by ATP-dependent pyrophosphorylation of 6-hydroxymethyl-7,8-dihydropterin (6-HMDP, **Figure 1A**). The resulting product is combined with *p-*ABA by DHPS to yield 7,8-dihydropteroate (7,8-DHP, **Figure 1A**). To produce THF, 7,8-dihydropteroate undergoes consecutive conjugations with glutamic acid followed by reduction of the pteridine core (Cossins and Chen, 1997).

**Figure 1.**
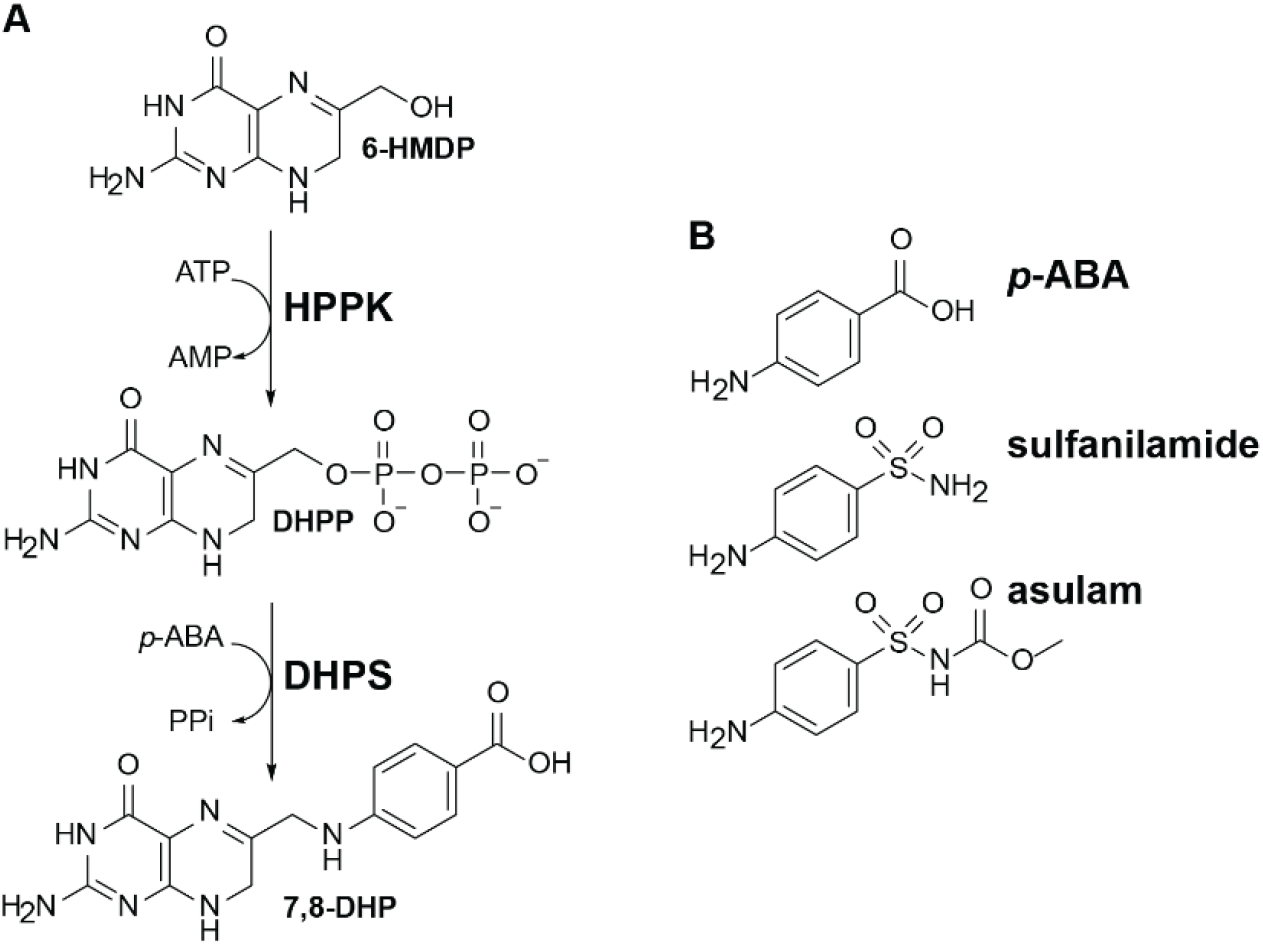
HPPK/DHPS is a herbicide target for sulfonamides that mimic DHPS substrate *p*-ABA. **(A)** Schematic of HPPK/DHPS mechanism. HPPK catalyses ATP-mediated pyrophosphorylation of 6-hydroxymethyl-7,8-dihydropterin (6-HMDP) to yield DHPP. DHPS then adds *para*-aminobenzoic acid (*p*-ABA) to DHPP forming 7,8-dihydropteroate (7,8-DHP). **(B)** DHPS substrate *p*-ABA and its antibiotic and herbicidal mimics sulfanilamide and asulam.

Although several of these steps in folate biosynthesis are targeted by antibacterials, antimalarials or human chemotherapies, asulam is the only herbicide targeting folate biosynthesis in plants. Asulam is a sulfonamide like the antibacterial ‘sulfa’ drug sulfanilamide (**Figure 1B**). Asulam and other sulfa drugs mimic the DHPS substrate *p*-ABA and inhibit enzyme activity by forming inactive adducts with 6-HMDP that inhibit downstream steps in folate synthesis (Roland et al., 1979; Chakraborty et al., 2013). Although popular for treating bacterial infections and malaria (Fernández-Villa et al., 2019), resistance to them is seen with mutations to residues capping the DHPS active site, making sulfa drugs that protrude beyond the DHPS surface more susceptible to resistance (Yun et al., 2012; Pornthanakasem et al., 2016; Griffith et al., 2018). Despite the similar structure of sulfa drugs and asulam, no equivalent resistance mutations in plant DHPS have been reported to date (Heap, 2021).

Although most bacteria have separate genes for HPPK and DHPS, in plants they are expressed from a single gene to give a conjoined, bifunctional enzyme, with the N-terminal HPPK connected by a short linker region to DHPS (Prabhu et al., 1997; Rébeillé et al., 1997). Like plants, protozoans including *Toxoplasma gondii* (Pashley et al., 1997) and *Plasmodium* spp. (Triglia and Cowman, 1994) also have bifunctional HPPK/DHPS. Some bacteria are unusual by having bifunctional HPPK/DHPS e.g. *Francisella tularensis* (Pemble et al., 2010) or a bifunctional dihydroneopterin aldolase (DHNA)-HPPK like *Streptococcus pneumoniae* where DHNA catalyses formation of the HPPK substrate (Arnaud et al., 2006). *Saccharomyces cerevisiae* expresses a trifunctional DHNA-HPPK/DHPS (Ulrich et al., 2004; Lawrence et al., 2005).

Although the *HPPK/DHPS* gene is widespread in plants, its protein product has been biochemically characterised only in *Pisum sativum* (Rébeillé et al., 1997) and *Arabidopsis thaliana* (Prabhu et al., 1997; Storozhenko et al., 2007). Some plants such as pea (Rébeillé et al., 1997) and wheat (McIntosh et al., 2008) only have a mitochondrial *HPPK/DHPS* gene, whereas *A. thaliana* has two 81% identical genes encoding a mitochondrial (mitHPPK/DHPS, At4g30000) and a cytosolic (cytHPPK/DHPS, At1g69190) form (Storozhenko et al., 2007). MitHPPK/DHPS is directly involved in folate biosynthesis and is essential, whereas the non-essential cytHPPK/DHPS has been shown to play a role in seed germination and stress response by an unknown mechanism that is apparently folate-independent (Navarrete et al., 2012). Although non-essential, the *A. thaliana* cytHPPK/DHPS gene rescued a yeast mutant devoid of HPPK/DHPS demonstrating *in vivo* that it has the catalytic capability of a mitochondrial enzyme and is suggestive that it has a catalytic role in the cytosol (Storozhenko et al., 2007). Current mechanistic insights into HPPK and DHPS catalysis rely on crystal structures of these enzymes from bacteria, fungi and protozoa. Whether individually expressed or as a part of multicomponent enzymes, HPPK and DHPS enzymes are structurally conserved across species with a high sequence similarity of active site residues (**Figure 2**).

**Figure 2.**
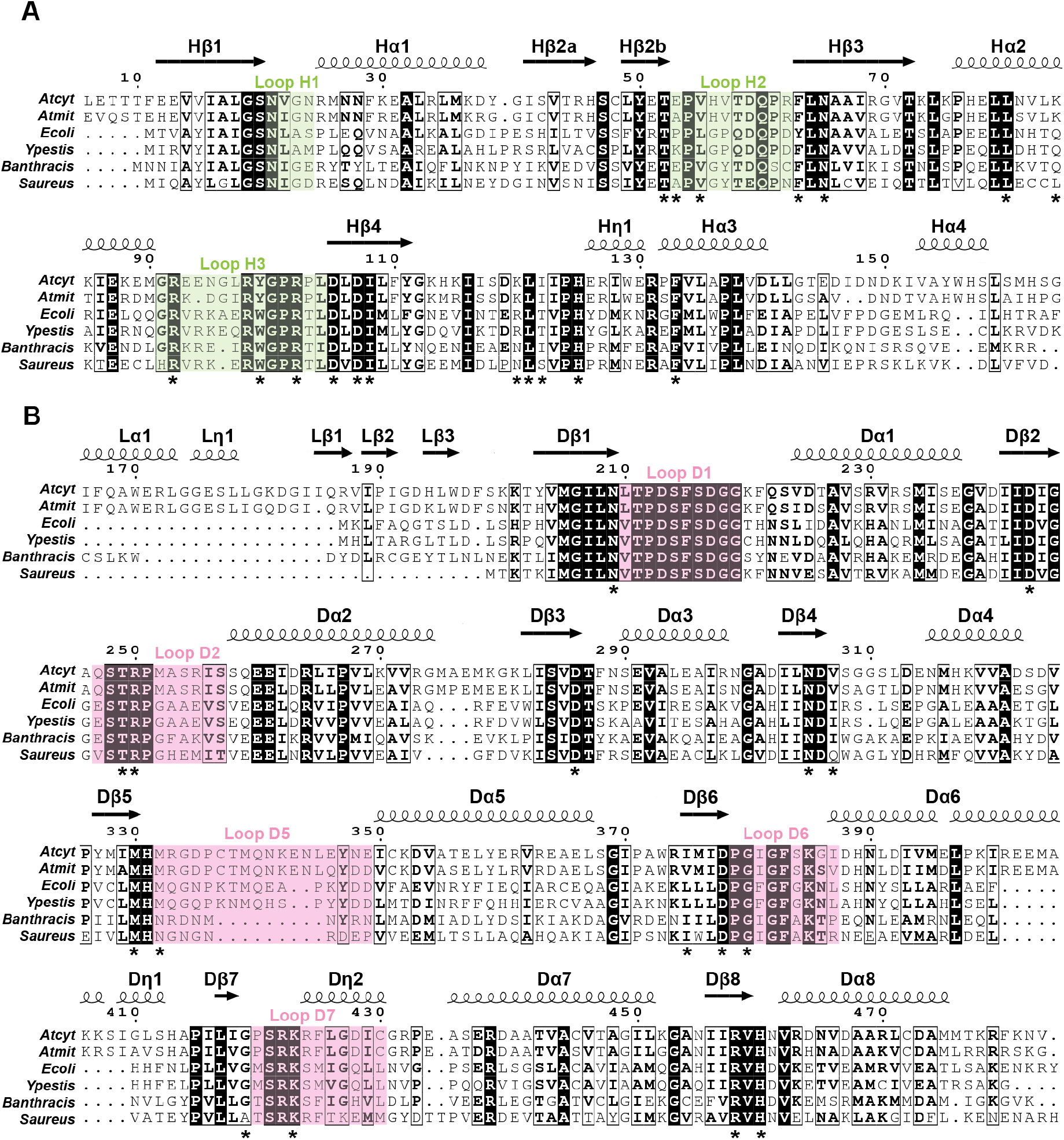
The HPPK/DHPS active site residues are conserved. Alignment of *A. thaliana* cytosolic and mitochondrial forms of **(A)** HPPK and **(B)** DHPS domains with microbial sequences. Identical (white font) and similar residues (bold font) are noted. Catalytic loops H1-H3 in HPPK are highlighted green whereas DHPS catalytic loops D1-D7 are pink. Residues involved in catalysis are asterisked. β-strands (marked as arrows), α-helices and 3_10_ helices (marked as coils) are annotated by the symbols β, α and η, respectively.

Given the similarity of asulam to antibacterial sulfonamide drugs, to develop novel DHPS inhibitors as herbicides it is important to contrast the structures of the plant enzyme against their bacterial homologs. To better understand the interaction of DHPS and asulam we solved the crystal structure for *A. thaliana* cytHPPK/DHPS at a resolution of 2.6 Å. We found both components are similar to their microbial counterparts with an overall structural root mean square deviation (r.m.s.d.) of 1.1-2.3 Å in C_α_ atoms. We used this structure to model *A. thaliana* mitHPPK/DHPS, which we were unable to express in a soluble form. We also compared the herbicidal and antibiotic activities of seven sulfonamides (including asulam) to understand which had cross-kingdom efficacy. Overall, the structural data presented provide insight to help protect this mode of action and highlights HPPK as a new potential herbicide target in the class of folate biosynthesis inhibitors.

## Results and Discussion

### The HPPK/DHPS active site is conserved across kingdoms

Comparing *A. thaliana* HPPK/DHPS with sequences of microbial species shows conservation of the catalytically relevant HPPK and DHPS regions (**Figure 2, Supplemental Figure 1**). The DHPS domains of bifunctional HPPK/DHPS from *S. cerevisiae* and *Plasmodium* species contain long insertions (**Supplemental Figure 1**). In contrast, the cytDHPS and mitDHPS are closer in length to bacterial DHPS enzymes and share 28-43% identity (**Table S1**). Secondary structure is conserved between plants and microbes, but the linker region varies in length for microbial species with a bifunctional HPPK/DHPS (**Figure 2, Supplemental Figure 1**).

Alignments of HPPK/DHPS from throughout the plant kingdom reflect active site conservation with sequence identity of 47-81% (**Supplemental Figure 2, Table S2**). For agricultural relevance, we included weed species commonly treated with asulam, including the grasses *Lolium multiflorum, Echinochloa crus-galii, Cyperus rotundus* and *Poa annua* (**Table S2**). Collectively, the sequence alignments reveal that the HPPK/DHPS active sites are not only conserved within the plant kingdom but across microbial species, as well.

### Crystal structure of *A. thaliana* cytosolic HPPK/DHPS

An N-terminally 6-His tagged *A. thaliana* cytHPPK/DHPS expressed in *E. coli* was crystallised and its structure determined by molecular replacement in an unliganded form. Attempts to co-crystallise cytHPPK/DHPS with asulam only gave crystals whose diffraction lacked electron density for the herbicide, suggesting pterin binding might be required to favour binding of asulam within the *p-*ABA binding pocket.

Crystals of cytHPPK/DHPS diffracted to a resolution of 2.6 Å and belong to the space group *C* 2 2 2_1_ with two protein molecules per asymmetric unit (**Table 1, Figure 3A-B**). Each monomer consists of an N-terminal HPPK domain (residues 1-160) linked to the C-terminal DHPS domain (residues 203-483) by a structured linker region (residues 161-202). Inspection of the asymmetric unit revealed that the two monomers do not form a dimer. The interaction between the monomers occurs between HPPK catalytic loops H1, H2 and H3 (**Figure 2, Figure 3B**), which is in contrast to bifunctional HPPK/DHPS structures for *P. falciparum* and *P. vivax* that interact at their DHPS domains (Lawrence et al., 2005; Yogavel et al., 2018; Chitnumsub et al., 2020). However, each cytHPPK/DHPS monomer within the asymmetric unit forms a homotypic dimer with the same monomer from a neighbouring asymmetric unit (A:A’ and B:B’ rather than A:B). The dimerisation interface is via the DHPS domain of each monomer (**Figure 3C**) through helices Dα6, Dη3, Dα7 and Dα8. Assignment of these interaction interfaces is supported by the PDBePISA server (Krissinel and Henrick, 2007) which does not identify the A:B interaction as a probable interface (Δ^i^G p-value 0.66), while the A:A’ (*−x, y, −z+1/2*; **Figure 3C**), or B:B’ (*−x, y, −z−1/2*, **Supplemental Figure 3A**) interfaces are strongly predicted as real (Δ^i^G p-values <0.01). Furthermore, the A:A’ and B:B’ dimers are very similar (r.m.s.d. 1.2 Å for 423 pairs of C_α_ atoms), and the observed arrangement resembles previously determined HPPK/DHPS dimer structures (**Supplemental Figure 3B**) (Lawrence et al., 2005; Yogavel et al., 2018; Chitnumsub et al., 2020). In summary, the asymmetric unit contains two halves of two crystallographic dimers.

**Table 1.**
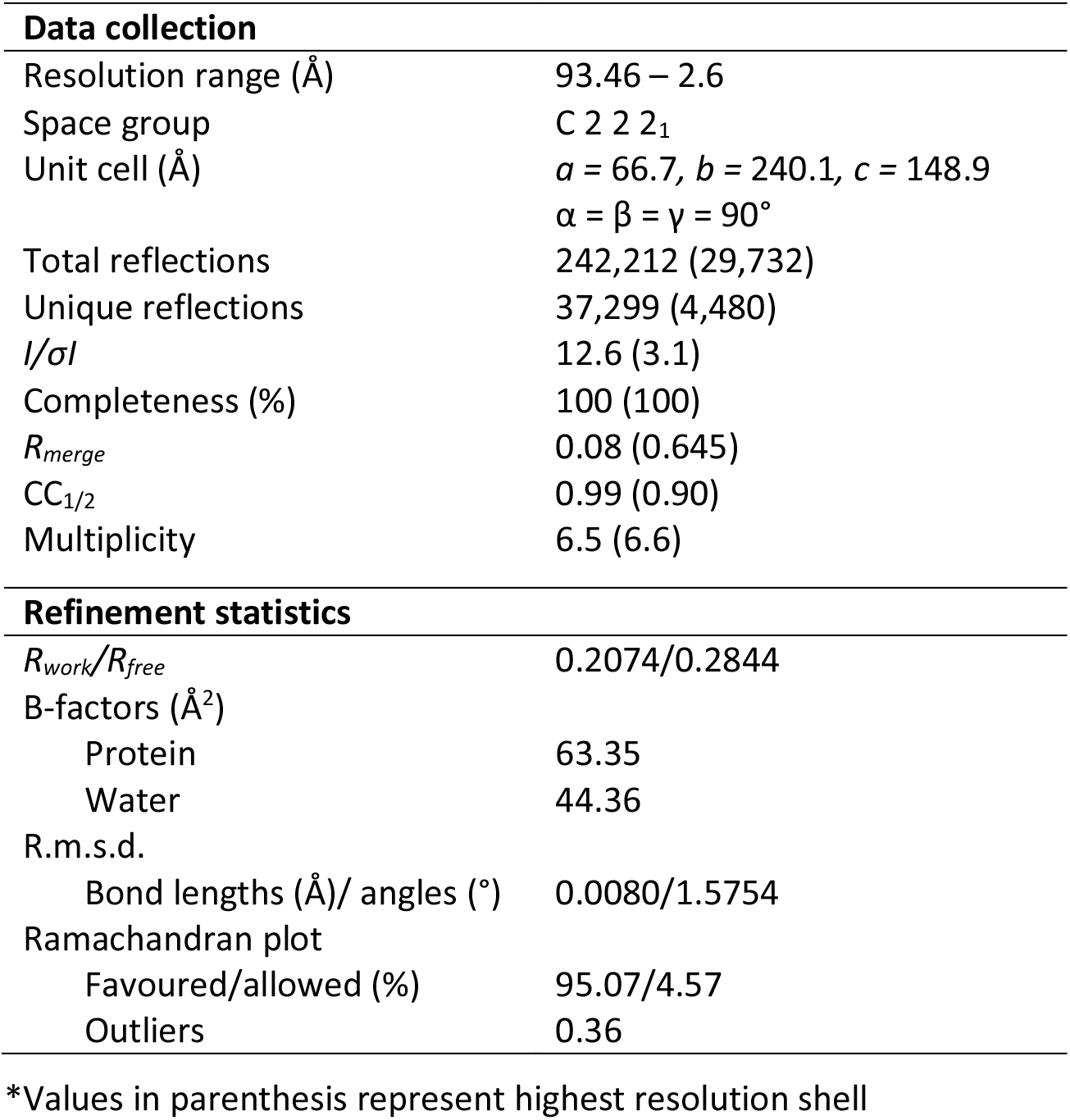
Crystallographic statistics for *A. thaliana* cytHPPK/DHPS.

**Figure 3.**
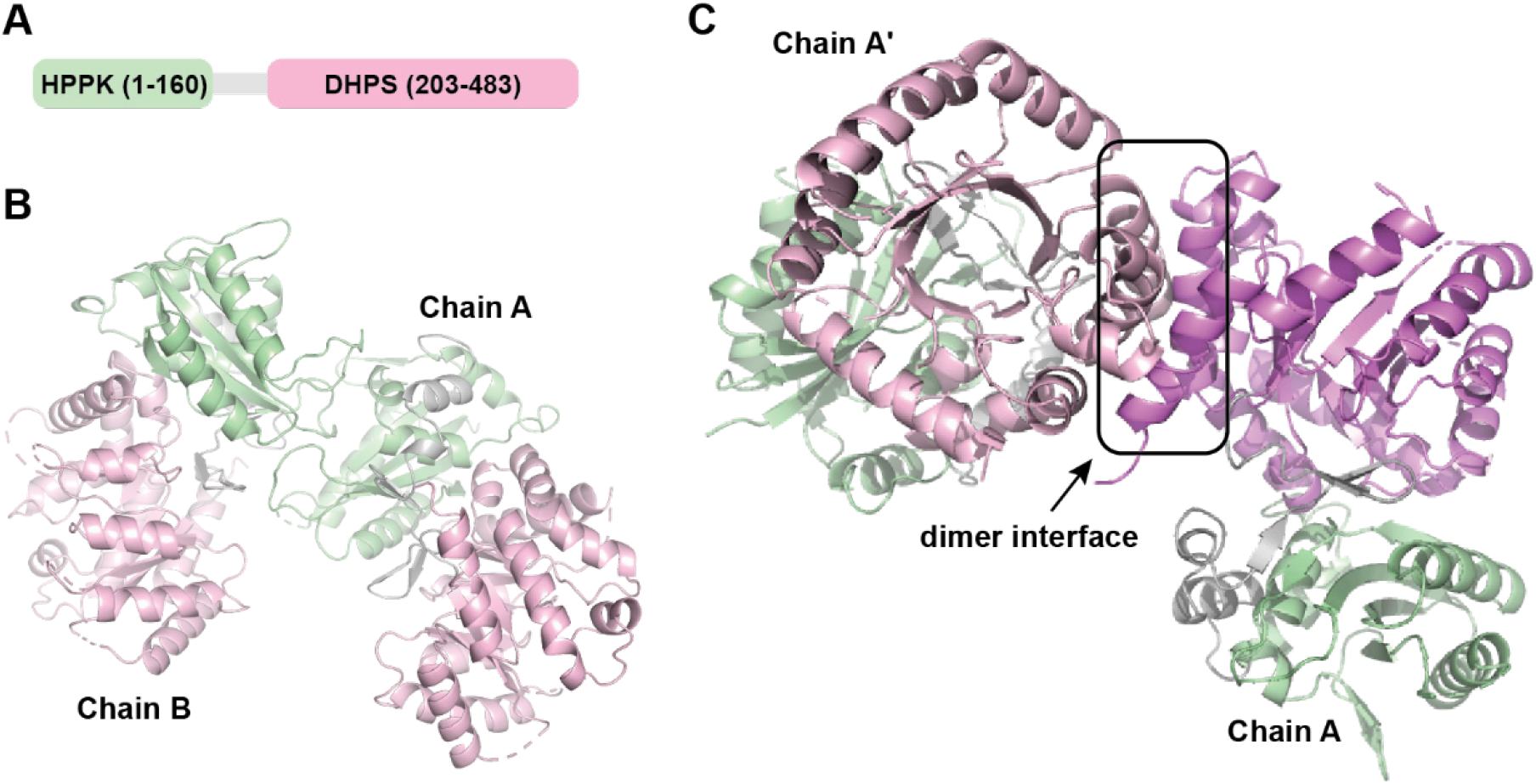
*A. thaliana* cytHPPK/DHPS forms a crystallographic dimer. **(A)** Simplified schematic of cytHPPK/DHPS. **(B)** Monomers in the crystallographic asymmetric unit interact at their HPPK (green) domain, which is connected to DHPS (pink) by a structured linker (grey). **(C)** Crystallographic dimer between chain A of the asymmetric unit and its symmetry mate A’ with an interface comprising Dα6, Dη3, Dα7 and Dα8.

The two molecules in the asymmetric unit differ mainly in their flexible loop regions (r.m.s.d. of 1.2 Å in C_α_ atoms). Additionally, disordered loops account for one region missing electron density in HPPK (chain A - loop H3 residues 93-100) and four regions of DHPS (chain A - loop D1 residues 213-222, loop D2 residues 247-256, residues 309-312, loop D5 residues 333-348; chain B - loop D1 residues 213-221, loop D2 residues 247-259, residues 311-313, loop D5 residues 333-347). Due to the flexibility of the catalytic loops in HPPK and DHPS, they are often disordered (Babaoglu et al., 2004; Lawrence et al., 2005; Pemble et al., 2010), as reflected by missing loops within the structure.

Like its microbial homologs, cytHPPK adopts an αβα ferredoxin-like fold comprised of a central, four-strand antiparallel β-sheet (Hβ2-Hβ3-Hβ1-Hβ4) sandwiched between four α-helices (Hα1-Hα2 on one face, and Hη1-Hα3-Hα4 on the other face) (**Figure 2, Figure 4A**). In contrast to monofunctional HPPKs, the 42-residue linker of cytHPPK/DHPS begins with two successive helices (Lα1 and Lη1) that stabilise the HPPK domain, followed by a short 3-residue β-strand (Lβ1) that associates with Hβ3 in HPPK, and terminates in a β-hairpin cap (Lβ2-turn-Lβ3) that closes over the N-terminal end of the TIM-barrel in DHPS (**Figure 2, Figure 3B**). The cytDHPS structure has a typical TIM-barrel like fold (Babaoglu et al., 2004; Lawrence et al., 2005) comprised of an eight-stranded β-barrel surrounded by eight α-helices (**Figure 2, Figure 5A**). The β-sheet appearance of residues Ile303 to Asp306 in what should be Dβ4 is not defined in the chain B structure likely due to the disordered loop connecting this segment to Dα4.

**Figure 4.**
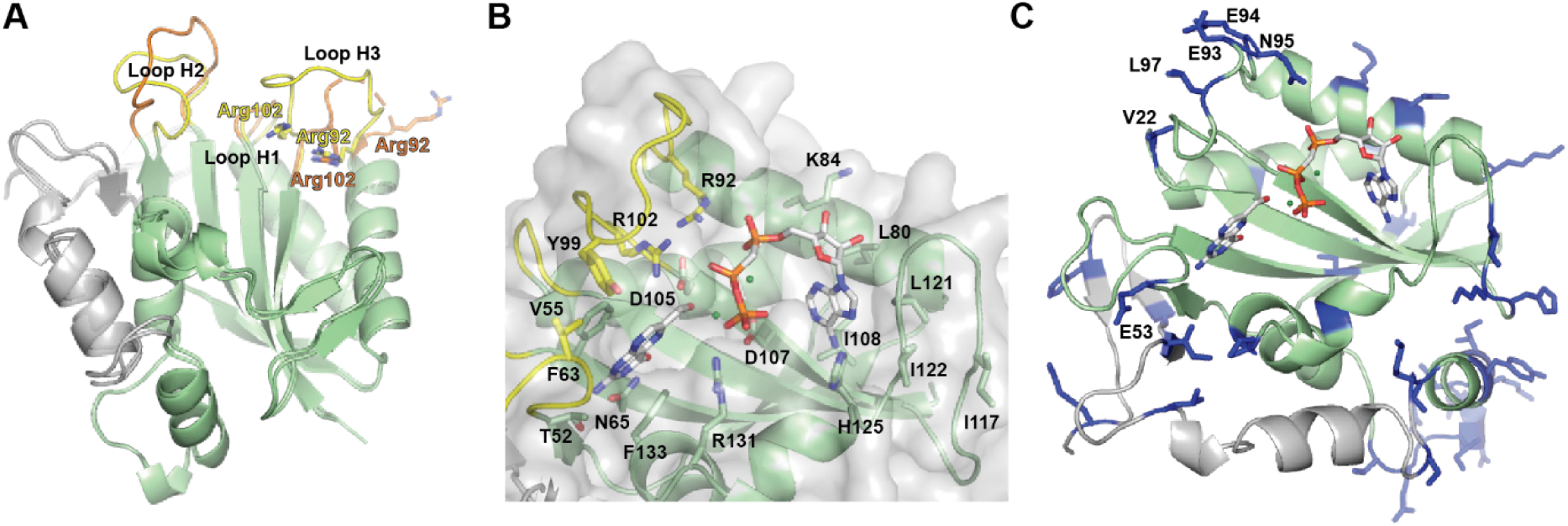
cytHPPK has a conserved active site flanked by catalytic loops in different conformations. **(A)** Superposition of chain A and chain B cytHPPK domains, with catalytic loops 1 to 3 coloured yellow (chain A) or orange (chain B). Disordered regions in loop H3 (chain A) are indicated by a dashed line. **(B)** HPPK substrates 6-HMDP and the non-hydrolysable ATP-mimic diphosphomethylphosphonic acid adenosyl ester (AMPcPP) from the *F. tularensis* structure (PDB ID 3mco) are superposed in cytHPPK with catalytically relevant residues shown as sticks. Ligand atoms are shown in grey (carbon), blue (nitrogen), red (oxygen), orange (phosphorus) and green (magnesium), respectively. **(C)** Residues that differ between mitHPPK and cytHPPK are highlighted (dark blue sticks) showing general active site conservation between homologs, with a few labelled exceptions.

**Figure 5.**
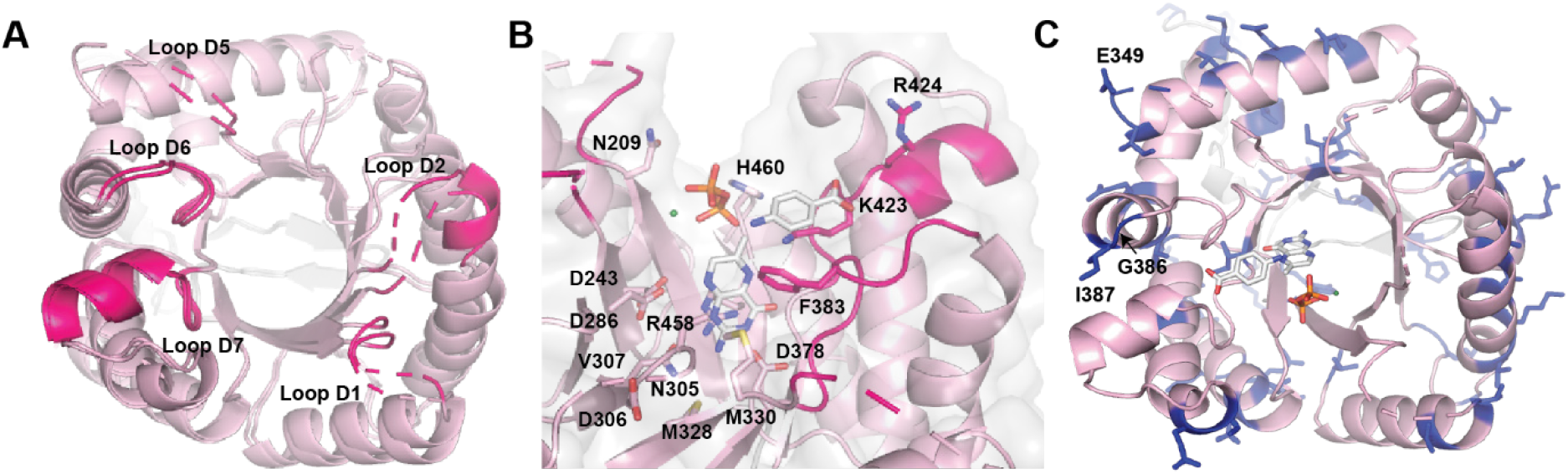
cytDHPS is structurally conserved with microbial counterparts. **(A)** Superposition of cytDHPS domains from chain A and chain B shows both monomers adopt similar conformations. Catalytic loops 1, 2, 5, 6 and 7 are highlighted dark pink, with regions of disorder indicated by dashed lines. **(B)** DHPS substrates DHP+, PPi and *p*-ABA from *Y. pestis* (PDB ID 3tyz) were superposed to show catalytically relevant residues (sticks). Ligand atoms are shown in grey (carbon), blue (nitrogen), red (oxygen), orange (phosphorus) and green (magnesium), respectively. **(C)** Residues non-identical to mitHPPK are highlighted dark blue, revealing active site conservation apart from Glu346 (in disordered loop D5, not observed) and Gly386 (labelled).

In *P. falciparum* HPPK/DHPS, a patch of positively charged residues adorn the surface between HPPK and DHPS active sites and are hypothesised to facilitate channelling of the negatively charged DHPP from one site to the other (**Supplemental Figure 4**) (Chitnumsub et al., 2020). In some microbial parasites, another bifunctional folate biosynthesis enzyme was hypothesised to channel substrate by a path of positively charged surface residues connecting the active sites of the two domains (Anderson, 2017). Here, for cytHPPK/DHPS and a model of mitHPPK/DHPS generated using the cytHPPK/DHPS crystal structure as a template using iTASSER (Yang et al., 2015a), we found the domains were connected by a largely electronegative stretch of residues (**Supplemental Figure 4**), making the hypothesis of substrate channelling unlikely. It is worth noting that *P. falciparum* and *P. vivax* have multiple long insertions within their HPPK and DHPS domains that alter surface architecture (**Supplemental Figure 1**) (Yogavel et al., 2018; Chitnumsub et al., 2020). In contrast, the *A. thaliana* enzymes are closer in length and sequence identity to bacterial monofunctional enzymes (**Figure 2, Supplemental Table 1**). Enzyme kinetics of *P. sativum* HPPK/DHPS showed catalysis of 6-HMDP by HPPK was rate-limiting for the production of 7,8-DHP by the bifunctional enzyme (Mouillon et al., 2002). In the absence of *p*-ABA, the product of HPPK, DHPP accumulated in the medium, but was rapidly turned over upon addition of *p*-ABA (Mouillon et al., 2002). The authors reasoned that substrate channelling seemed less likely given the propensity of DHPP to freely equilibrate with the external media, and that its rapid clearance by DHPS was due to high specific activity (Mouillon et al., 2002). Taken together with our structural data, this suggests that substrate channelling does not play a role in HPPK/DHPS activity in *A. thaliana*.

### The cytHPPK catalytic loops adopt different conformations between monomers

Superposition of individual monomers within the cytHPPK/DHPS asymmetric unit revealed differences in the catalytic loops (**Figure 4A**). HPPK catalysis requires participation from residues in loops H1, H2 and H3 (**Figure 2, Figure 4A**) that delineate the enzyme active site. In cytHPPK, loop H1 adopts similar conformations in each chain. In chain B, loop H2 projects outwards relative to chain A and loop H3 projects towards the active site, showing clear electron density for key catalytic residues Arg92 and Arg102. Chain A loop H2 is angled slightly inwards, and eight residues in chain A loop H3 are disordered, with Arg92 removed from the active site and Arg102 positioned within the active site (**Figure 4A**).

To examine active site architecture, cytHPPK chain B was used as clear electron density was observed for the entire domain. Enzyme substrates 6-HMDP and the non-hydrolysable ATP-mimic AMPcPP were superposed onto the cytHPPK site based on structural similarity to the *F. tularensis* enzyme (r.m.s.d. of 1.4 Å in C_α_ atoms) (**Figure 4B**) (Pemble et al., 2010). The binding site for AMPcPP is situated closer to the surface, where residues Leu80, Lys84, Ile108, Leu121, Ile122 and His125 could stabilise the pocket around the adenosine ring of ATP. The phosphate groups are proposed to be further stabilised by electrostatic interactions with residues Lys84, Arg92, Arg102, His125 and Arg131 (Pemble et al., 2010). As the cytHPPK binding site is unliganded, Lys84 and His125 are moved slightly away from the active site. The essential Mg^+2^ cofactors are coordinated by two conserved aspartate residues, Asp105 and Asp107, however in cytHPPK, Asp105 is pointed away from Asp107 in the absence of Mg^+2^ atoms. In *Ft*HPPK, this Mg^+2^ atom can be seen to coordinate the α- and β-phosphates of AMPcPP, and the second Mg^+2^ atom coordinates the β- and γ-phosphate and orients the 6-hydroxymethyl group of 6-HMDP towards the pyrophosphate moiety of AMPcPP for transfer (Pemble et al., 2010). At the base of the pterin-binding pocket, Phe63 and Phe133 are clearly positioned in a highly conserved π-π stacking interaction where they could flank the pterin ring of 6-HMDP. Hydrogen bonding interactions between Thr52 and the conserved Asn65 tether 6-HMDP to the base of the pocket by its nitrogen face, and Val55 and Tyr99 complete the binding pocket.

HPPK catalysis is proposed to involve six steps beginning with Mg^+2^-ATP binding to HPPK (Blaszczyk et al., 2000). Then, outward movement of loop H3 (by ~20 Å) triggers key arginine residues (Arg92 and Arg102 in cytHPPK) to complete the active site. Next, 6-HMDP binds and loops H1 and H3 close around it, with a tryptophan residue (Tyr99 in cytHPPK) moving in to seal the active site closed. After the reaction is completed, loop H3 moves back outwards to release the product DHPP.

Based on the currently known reaction trajectory of HPPK catalysis (Blaszczyk et al., 2000), it would appear that chain A in unliganded cytHPPK adopts a conformation either poised to accept 6-HMDP or to release the product DHPP. Chain B, on the other hand reflects a conformation facilitating catalysis within the active site, as loop H3 is moved inwards, with both Arg92 and Arg102 facing the substrate-binding pockets and Tyr99 partially sealing the active site.

To investigate how the sequence differences in the mitochondrial HPPK/DHPS of *A. thaliana* might influence HPPK catalysis, we mapped the different residues onto the cytHPPK structure (highlighted in blue, **Figure 4C**). Within catalytically relevant loop regions, cytHPPK differs from mitHPPK at six amino acid residues. In cytHPPK, loop H1 residue Val22 (Ile97 in mitHPPK) and loop H3 residue Leu97 (Ile171 in mitHPPK) are similar in size and hydrophobicity. Interestingly, where cytHPPK has Glu53 in loop H2, there is an Ala128 in mitHPPK. Whether a larger, negatively charged residue affects the ability of loop H2 to stabilise the active site pocket in HPPK is unclear. Although most microbial HPPK enzymes have a charged residue at this site, *E. coli* HPPK has a proline and *S. aureus* an alanine, suggesting glutamate at this position might not adversely affect enzyme catalysis. In loop H3 of cytHPPK, Glu93 (Lys168 in mitHPPK) is followed by Glu94 and Asn95 (Asp169 in mitHPPK). Considering the important role of loop H3 in mediating product binding and release, it is possible that the presence of Glu93-Glu94-Asn95 in cytHPPK affects its interactions with substrate/product sufficiently to affect its activity. Aside from these residues in the catalytic loops, variations between the mitHPPK and cytHPPK largely decorate surfaces away from the active site.

### The cytDHPS active site is structurally conserved

In contrast to the differences between HPPK monomers, superposition of the DHPS crystallographic chains A and B revealed only subtle differences (**Figure 5A**). Loops D1, D2, D4 and D5 were disordered in both crystallographic monomers, and are often disordered in DHPS crystal structures. Among the loops connecting individual β-sheets to α-helices in the TIM-barrel, loops D1, D2, D5, D6 and D7 are actively involved in mediating catalysis or stabilising the active site.

To understand the *A. thaliana* DHPS structure in context of its bound substrates, we superposed DHP+, pyrophosphate and *p*-ABA into the cytDHPS active site (**Figure 5B**) based on a crystal structure of the substrate-bound DHPS from *Yersinia pestis* (r.m.s.d. of 1.2 Å in C_α_ atoms) (Yun et al., 2012). When crystals of *Y. pestis* DHPS were soaked in DHPP and *p*-ABA, the pyrophosphoryl group from DHPP was cleaved and remained in the active site (Yun et al., 2012). Comparison with microbial DHPS shows these two binding pockets in cytDHPS – a pterin-binding site at the opening of the β-barrel, and a *p*-ABA binding site closer to the surface formed by loops D1, D2 and D7.

The pterin-binding pocket contains two aspartate residues (Asp286 and Asp378) proposed to stabilise the resonance forms of the pterin-ring during pyrophosphoryl cleavage, with the latter residue essential for DHPP binding (Yun et al., 2012). In addition to Asp286 and Asp378, Asn305 and Lys423 would form a part of the hydrogen bond network stabilising the pterin ring (Yun et al., 2012). In the unliganded cytDHPS, Arg458 overlaps the pterin binding site – in a substrate-bound structure, this arginine would probably engage in π-stacking interactions parallel to the pterin ring such that it could also interact with the pyrophosphate moiety. The superposed pyrophosphate group sits in an adjacent cleft comprised of Ser214, Ser216 and Asp217 (from loop D1), Ser248 and Thr249 (from loop D2), and Arg458 and His460 which form the anion-binding pocket (Yun et al., 2012). At this site, a Mg^+2^ ion would form part of a coordinated network between the pyrophosphate oxygen atoms, Asn209 and water molecules. This Mg^+2^ ion is proposed to assist pyrophosphate release from the DHPS active site, as well as stabilise the substructure formed by catalytic loops D1 and D2 (Yun et al., 2012; Bourne, 2014).

Key interacting residues in the *p*-ABA binding pocket include Phe215 (loop D1), Pro251 (loop D2), Phe383 (loop D6) and Lys423 (loop D7). In *Yp*DHPS, the *p*-ABA carboxylate group hydrogen bonds to a neighbouring serine residue in loop D7 (Arg424 in cytDHPS) and is stabilised by the helix dipole of Dα7 (Yun et al., 2012). In cytDHPS, the involvement of functionally critical loops D1 and D2 in *p*-ABA binding cannot be seen due to loop disorder, however, Phe383, Lys423 and Arg424 can be seen to adopt orientations similarly to their counterparts in *Yp*DHPS.

Drawing from a series of bacterial DHPS structures crystallised with various substrate/products, a mechanism was proposed for 7,8-DHP synthesis (Babaoglu et al., 2004; Yun et al., 2012). The reaction begins with DHPP binding, which stabilises mobile catalytic loops including loops D1 and D2 into forming the *p*-ABA binding pocket. Pyrophosphate cleavage from DHPP occurs first by an S_N_1 reaction, with both molecules retained in their respective binding pockets. Mg^+2^ facilitates departure of the pyrophosphate from the active site, allowing *p*-ABA to be covalently attached to the unphosphorylated pterin to yield 7,8-DHP.

It is proposed that in the absence of pterin substrate, its binding site is occupied by an arginine residue in loop D2 (Arg250 in cytDHPS) where the guanidinium group of arginine coordinated with water molecules mimic the pterin substrate. This conformation was observed in native *B. anthracis* DHPS, where loop D2 also blocked access to the *p*-ABA binding site (Babaoglu et al., 2004). It is hypothesised that DHPP entry displaces the Arg-containing loop D2 which now forms the *p*-ABA binding pocket. Following product formation, this loop is suggested to swing back in to displace product from the active site. This is predicted to be a mechanism to prevent *p*-ABA binding prior to pterin-binding as this would occlude the pterin-binding site and hinder catalysis (Babaoglu et al., 2004). This might explain our inability to crystallise DHPS with asulam alone. Considering that cytDHPS is unliganded, it could be expected that loop D2 would be moved in towards the pterin-binding pocket. However, several crystal structures of microbial DHPS enzymes also display disorder in loop D2 in their native and ligand-bound forms (Babaoglu et al., 2004; Morgan et al., 2011; Yun et al., 2012). This could be due to the sheer flexibility of the longer catalytic loops, and the associated difficulty of capturing a crystal structure of DHPS in a catalytically relevant conformation.

The DHPS active site and the base of the pterin-binding pocket in particular are almost absolutely conserved across species. As with *At*HPPK, we mapped on to the cytosolic DHPS structure those sites at which amino acid sequences differed from the mitochondrial enzyme (**Figure 5C**). Within the cytDHPS active site there is a Glu346 (Gln417 in mitDHPS) and Glu349 (Asp420 in mitDHPS) in loop D5 and a Gly386 (Ser457 in mitDHPS) and Ile387 (Val458 in mitDHPS) in loop D6. All variable residues were strictly either on the outside of the enzyme surface or facing away from the substrate-binding sites, hinting at the structural basis for why cytHPPK/DHPS can fully complement the function of its mitochondrial counterpart in *S. cerevisiae* (Rébeillé et al., 1997).

### Sulfonamides including asulam demonstrate cross-kingdom activity against *A. thaliana* and *E. coli*

Based on the sequence and structural similarity of cytDHPS to microbial DHPS, sulfonamides could be expected to have similar *in vivo* inhibition against plants and microbes. Accordingly, we contrasted the herbicidal activity of selected sulfonamides with varying R-groups against their antibacterial activity (**Figure 6, Table 2**). Our results show that asulam is indeed the most potent herbicide amongst tested sulfonamides, suppressing *A. thaliana* growth at 0.5 mM following treatment pre-emergence (**Figure 6**). Sulfacetamide and sulfamethoxazole, both commercially used antibacterials, also demonstrate reasonable potency, with growth inhibition at 2 mM and 4 mM, respectively (**Figure 6**). Conversely, although sulfamethoxazole strongly inhibits *E. coli* growth (minimum inhibitory concentration of 0.031 mM), inhibition by asulam and sulfacetamide is weaker at 0.125 mM and 0.25 mM, respectively (**Table 2**). Compared to sulfamethoxazole, sulfadoxine demonstrated weaker herbicidal and antibiotic activities (**Figure 6, Table 2**). Amongst the sulfonamides tested, sulfamethoxazole was the strongest inhibitor of the Gram-positive *B. subtilis* (MIC of 0.25mM), whereas asulam and sulfacetamide were not effective within the tested range. It is also interesting to note that altering an R-group from a ketone (as in sulfacetamide) to an ester (as in asulam) enhanced the herbicidal and antibacterial capability of this class of DHPS inhibitors (**Figure 6, Table 2**).

**Table 2.** Minimum inhibitory concentration for DHPS inhibitors.

| **Compound #** | **Compound name** | ***E. coli*** | ***B. subtilis*** |
| --- | --- | --- | --- |
| **1** | sulfanilamide | >2 | 2 |
| **2** | sulfacetamide | 0.25 | >2 |
| **3** | asulam | 0.125 | >2 |
| **4** | sulfamethoxazole | 0.031 | 0.25 |
| **5** | sulfadiazine | 0.062 | 1 |
| **6** | sulfamethazine | 0.5 | >2 |
| **7** | sulfadoxine | 0.25 | 2 |
|  | glyphosate | >2 | >2 |
|  | oryzalin | >2 | >2 |

**Figure 6.**
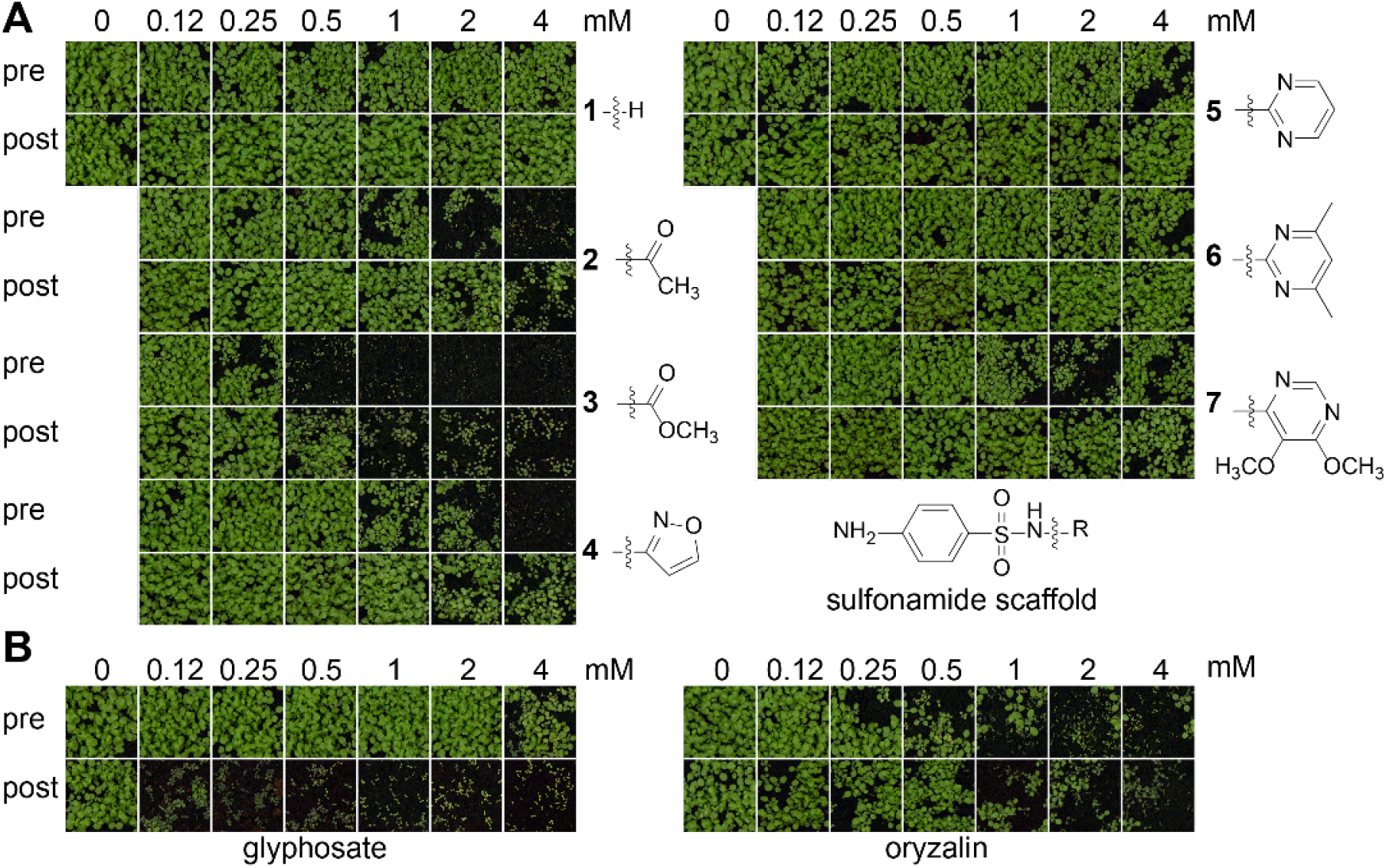
Asulam displays the strongest herbicidal activity against *A. thaliana* amongst a range of sulfonamides. Pre- and post-emergence herbicidal activities are shown for **(A)** sulfonamides (sulfanilamide **1**, sulfacetamide **2**, asulam **3**, sulfamethoxazole **4**, sulfadiazine **5**, sulfamethazine **6** and sulfadoxine **7**), and **(B)** herbicide controls glyphosate and oryzalin.

### Asulam has similar physicochemical properties to sulfonamide antibiotics

To investigate whether the lack of herbicidal activity by some sulfonamides could be a result of their physicochemical properties, we compared them to that of 359 commercial herbicides (**Figure 7**) (Sukhoverkov et al., 2021). The sulfonamides we tested largely cluster together within the expected range of commercial herbicides (**Figure 7**). Overall, the cluster analysis shows that sulfonamides including asulam have similar physicochemical properties that likely underpin their cross-kingdom activity and unlikely to be the basis for individual herbicidal potencies of asulam, sulfacetamide and sulfamethoxazole.

**Figure 7.**
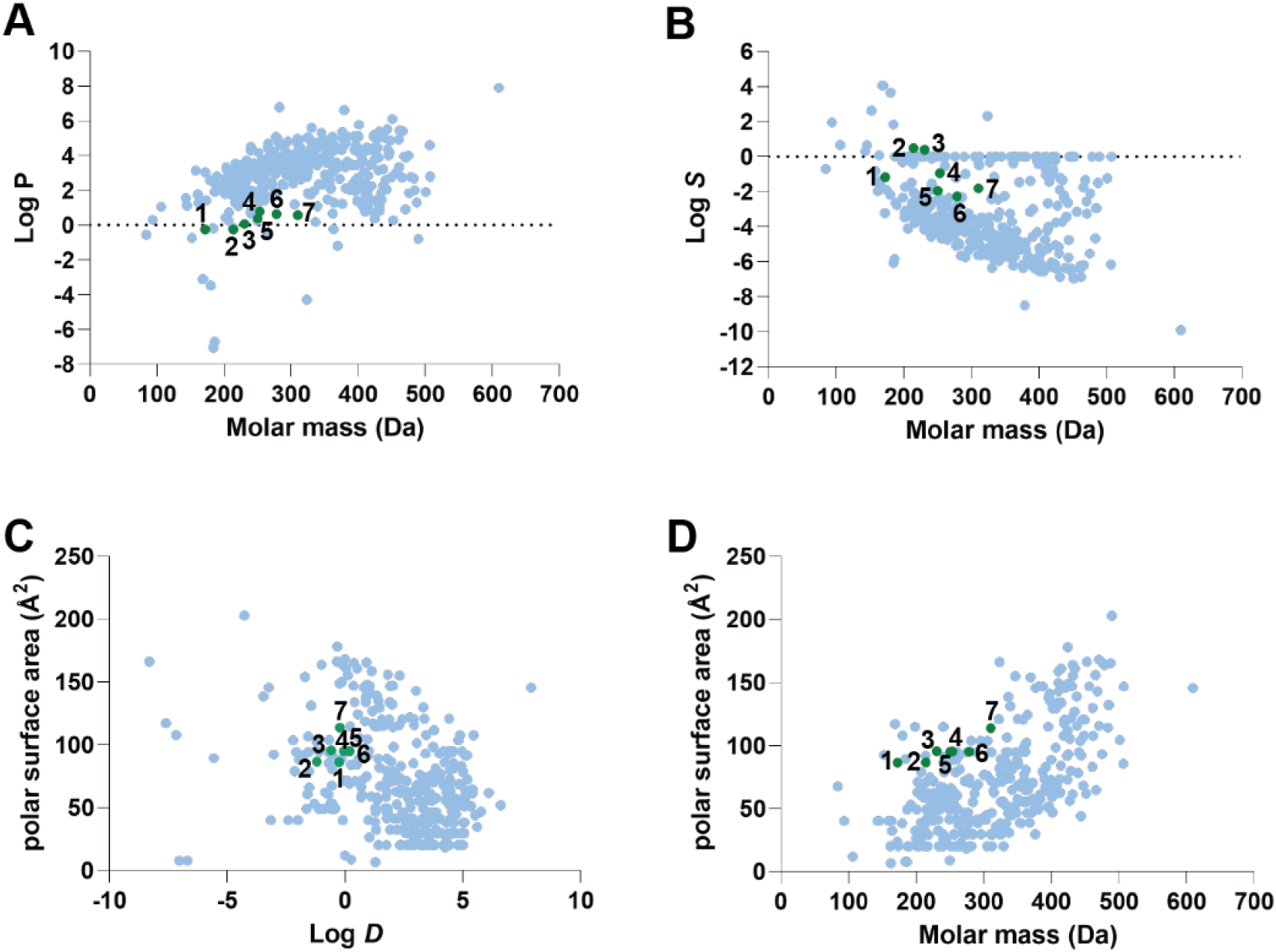
Physicochemical properties of sulfonamides and commercial herbicides. Physicochemical data for sulfonamides (green) were compared to 359 commercial herbicides (blue) (Sukhoverkov et al., 2021). Graphs compare **(A)** molar mass *vs* lipophilicity (Log *P*), **(B)** molar mass *vs* aqueous solubility (Log *S*), **(C)** distribution coefficient (Log *D*) *vs* polar surface area (Å^2^), **(D)** molar mass *vs* polar surface area (Å^2^).

### A model of mitDHPS and asulam shows the difficulty in developing plant-specific inhibitors

To study conservation of DHPS residues comprising the *p*-ABA/sulfonamide binding pocket, we examined key regions of catalytic loops D1, D2, D6 and D7 from 40 microbial species (**Supplemental Table 1**) and 40 plant species (**Supplemental Table 2**) using the WebLogo server (Crooks et al., 2004). These sequence logos illustrate which residues are conserved across plants and microbes (**Figure 8A**). To visualise the positioning of these residues relative to asulam, we created a mitDHPS homology model with SWISS-MODEL (Waterhouse et al., 2018) based on a *Y. pestis* DHPS holoenzyme crystal structure where the enzyme was captured in a catalytically active conformation (Yun et al., 2012). Using AutoDock Vina(Trott and Olson, 2010), the sulfonamides asulam and sulfamethoxazole were docked into the *p*-ABA binding pocket resulting in poses comparable to related structures, and predicted binding affinities of −5.7 and −6.5 kcal mol^−1^, respectively (**Figure 8B-C**).

**Figure 8.**
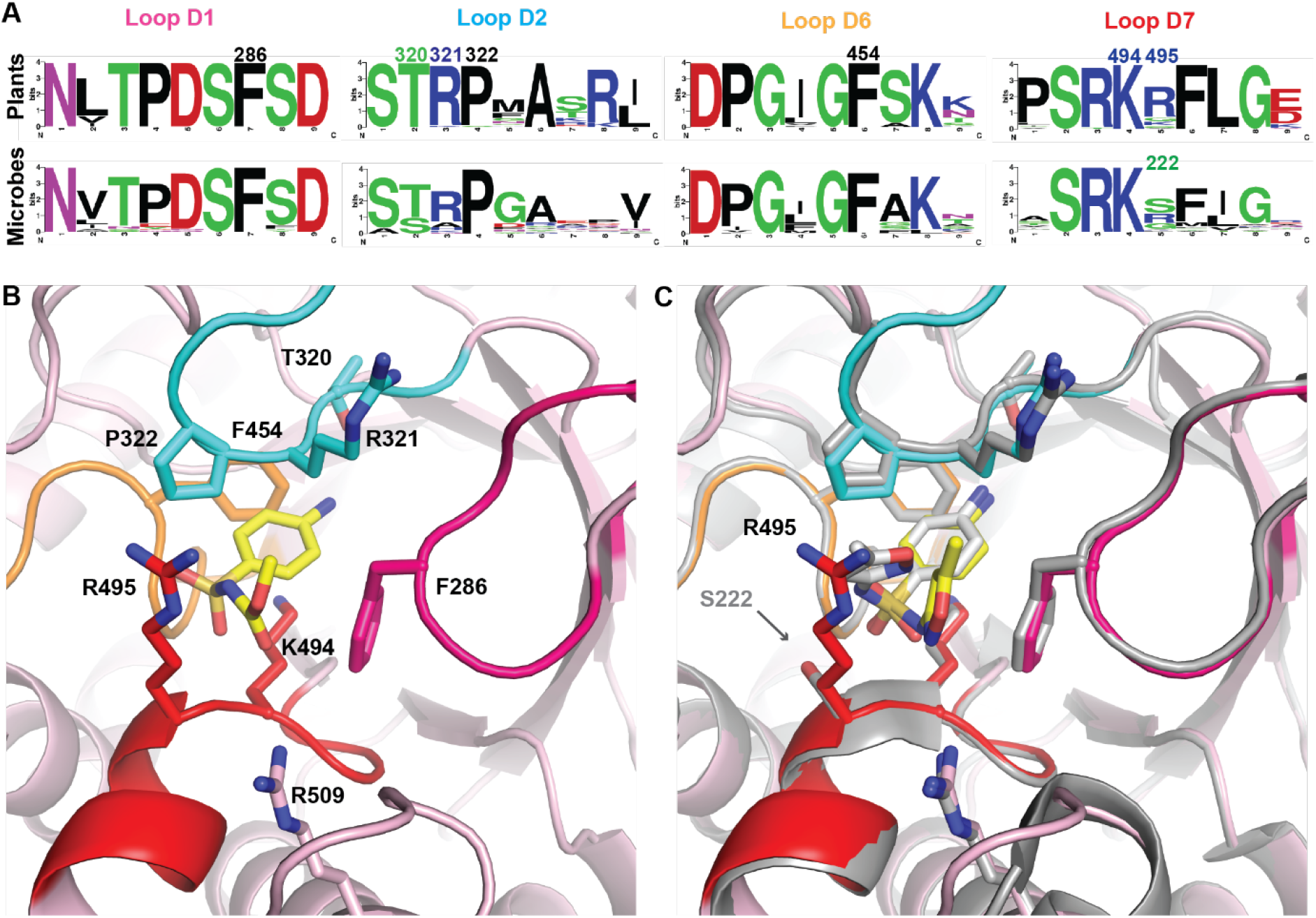
The sulfonamide binding pocket is conserved between plants and microbes. **(A)** Relative abundance of amino acid residues within subsections of DHPS catalytic loops D1, D2, D6 and D7 that form the *p*-ABA binding pocket. Contact residues that interact with *p*-ABA or sulfonamides are labelled with mitDHPS numbering. Arg509 falls within Dα7, adjacent to loop D7 **(B)** mitDHPS model with asulam docked in the *p*-ABA binding pocket within range of catalytic loop residues (colored as in panel A). Ligand atoms are shown in yellow (carbon), blue (nitrogen) and red (oxygen), respectively. **(C)** mitDHPS:sulfamethoxazole model superposed with a sulfamethoxazole-bound *Y. pestis* DHPS crystal structure (shown in grey; PDB ID: 3tzf) showing conservation of contact residues excepting Ser222 in *Yp*DHPS. Carbon atoms in sulfamethoxazole are shown in yellow (docked molecule) or grey (*Yp*DHPS-bound crystal structure).

The *p*-ABA binding pocket is formed by loops D1, D2, D6, D7 and the N-terminal region of Dα7. Key residues contributing to stabilizing *p*-ABA or sulfonamides at this site in mitDHPS would be Phe286 (loop D1); Thr320, Arg321 and Pro322 (loop D2); Gly453 and Phe454 (loop D6); and Lys494 and Arg495 (loop D7) (**Figure 8A-B**). Arg509 on Dα7 is within 3.5 Å of the docked sulfonamides, and would be expected to assist Arg495 in stabilizing the carboxylate moiety (or SO_2_ group in sulfonamides) by inference from the *Yp*DHPS structure bound to *p*-ABA or sulfamethoxazole (Yun et al., 2012). The catalytic loops in the mitDHPS model presented in **Figure 8** adopt a closed conformation reminiscent of the DHPS holoenzyme (Yun et al., 2012; Yogavel et al., 2018), whereas an unliganded *E*.*coli* DHPS crystal structure captures loop D1 in an open conformation (**Supplemental Figure 5**) (Achari et al., 1997).

Superposition of the sulfamethoxazole-bound *Yp*DHPS crystal structure to the mitDHPS:sulfamethoxazole model reveals absolute conservation of key residues excepting Ser221 in *Yp*DHPS at the site of Arg495 (**Figure 8C**). Although many plants have an arginine residue at this position, serine residues are also observed (**Figure 8A**), as in the weed *L. multiflorum* (**Supplemental Figure 2**). In contrast to the large, positively-charged Arg495, a serine residue at this position (as observed in *Y. pestis* and *E. coli*, see **Figure 8C, Supplemental Figure 5**) can potentially accommodate larger substituents, and may explain our observation that sulfadiazine was not herbicidal, but more antibacterial than asulam (**Figure 6, Table 2**). This position is a frequent site of mutation in microbial DHPS that results in sulfonamide resistance, which predominantly arises from mutations within loops D1, D2 and D7 (Yun et al., 2012; Yogavel et al., 2018; Chitnumsub et al., 2020). It has been noted that sulfa drugs with substituents extending beyond the Van der Waals surface of DHPS are more likely to select for resistance mutations, such as sulfadoxine whose bulky substituent moves catalytic loop D2 into a partially open conformation less able to stabilise inhibitor binding (Yun et al., 2012; Chitnumsub et al., 2020). Accordingly, lead compounds mimicking *p*-ABA must be designed to stay within the overall surface of DHPS and consider the role of loop D2 in mediating drug affinity as well as resistance. Interestingly in the sulfamethoxazole-bound *Yp*DHPS structure, the R-substituent still permits loop D2 to adopt a closed conformation covering the active site. Our findings that sulfamethoxazole, sulfacetamide and asulam demonstrated potent herbicidal and antibiotic activity could be a result of the similar size of their R-groups, which are smaller than sulfadoxine and sulfamethazine (**Figure 6**), and less likely to disrupt the stabilizing influence of loop D2. Failed attempts at docking sulfadoxine and sulfamethazine to mitDHPS may also be due to an inability to accommodate large conformational changes in loops D2 and D1 that may occur in the vicinity of their bulky R-groups.

### Exploring alternative HPPK/DHPS inhibitors as herbicides

Microbial drug discovery efforts have targeted the *p*-ABA and/or pterin-binding site of DHPS (Hammoudeh et al., 2014; Dennis et al., 2018), including pterin-like molecules pyrimido[4,5-*c*]pyridazines (Zhao et al., 2012) and derivatives of 5-nitroisocytosine (Babaoglu et al., 2004). Pterin-based DHPS inhibitors, though potent enzyme inhibitors *in vitro* (Zhao et al., 2012; Dennis et al., 2018) are often poor antibacterials *in vivo* (Lever et al., 1985; Lever et al., 1986). To determine whether pterin-based inhibitors have potential as herbicides, we synthesised a series of guanine-based inhibitors that mimic the pterin substrate of both HPPK and DHPS. Although compounds like these have promise as leads against microbial enzymes (Hevener et al., 2010; Dennis et al., 2014; Shaw et al., 2014; Dennis et al., 2018), we found them to have low solubility, which could be a contributing factor towards their complete lack of herbicidal and antibacterial activity (**Supplemental Figure 6, Supplemental Table 3**). However, analysis of the physicochemical properties of these pterins revealed that they are within the expected range of commercial herbicides (**Supplemental Figure 7**). Pterin-mimicking inhibitors are less likely to give rise to resistance due to the absolute conservation of residues at the base of the DHPS active site, so screening a library of suitably substituted pterin-mimics might uncover lead compounds providing plant-specificity and which might be optimised using the structural data presented here.

Another avenue for herbicide design could be to exploit the opportunities presented by two substrate-binding sites in each of HPPK and DHPS. Herbicides that occupy both substrate sites simultaneously in each individual enzyme could be developed. Indeed, bisubstrate pterin-sulfa conjugate inhibitors targeting DHPS have been shown to have antibacterial activity and bind to *Y. pestis* DHPS by occupying both pterin and *p*-ABA pockets (Zhao et al., 2016). They could be a viable solution to developing potent inhibitors less prone to resistance as the absolute conservation of the pterin-binding pocket in DHPS is less likely to tolerate resistance mutations, even if resistance is more frequent at the sulfonamide binding pocket. Success with HPPK inhibitors, including bisubstrate HPPK inhibitors that occupy both the pterin pocket and ATP pocket, has been achieved mostly *in vitro*, and is yet to reveal promising *in vivo* antibacterial candidates (Shi et al., 2012; Chhabra et al., 2013; Dennis et al., 2014).

In summary, our crystal structure of the bifunctional *A. thaliana* cytHPPK/DHPS shows that plant enzymes are structurally conserved compared to their microbial counterparts and provides a structural context for the herbicidal activity of asulam and the sulfonamide antibiotics sulfacetamide and sulfamethoxazole. Although studies have investigated the off-target and toxicity effects of asulam, such as its inhibition of mammalian sepiapterin reductase (Yang et al., 2015b), its cross-kingdom activity against soil microbiota is less well-studied (European Food Safety Authority et al., 2018). Sulfonamide antibiotics are used in the livestock industry and have been reported to have phytotoxic effects (Liu et al., 2009; Piotrowicz-Cieślak et al., 2010; Cheong et al., 2020). As we demonstrate the herbicidal potency of sulfamethoxazole against *A. thaliana*, a previous study showed it was herbicidal against duckweed (*Lemna gibba*) (Brain et al., 2008). The conservation of key sulfonamide-interacting residues within the DHPS active site suggests that achieving plant specificity for a *p-*ABA mimicking inhibitor might not be trivial. Exploiting differences in pharmacokinetic or delivery mechanisms between plants and microbes might be an effective way to develop greater plant-specificity. Although asulam resistance in weeds has not yet been reported, the combined inhibition of HPPK and DHPS could provide a means to pre-empt resistance.

## Materials and Methods

### Sequence alignments and analysis

HPPK/DHPS sequences were retrieved from NCBI Protein BLAST using *A. thaliana* mitHPPK/DHPS as the query. Transcriptomes of weed species commonly targeted by asulam in the field were assembled using CLC Genomics Workbench v20.0.3 (Qiagen Aarhus A/S) from data in the NCBI Sequence Read Archive: *L. multiflorum* (accession SRR1648407)(Czaban et al., 2015), *E. crus-galli* (SRR8633067) (Fang et al., 2019), *C. rotundus* (SRR12887711) (Ji et al., 2021) and *P. annua* (SRR1633980) (Chen et al., 2016). Protein sequences of HPPK-DHPS from these species were obtained using tBLASTn by searching the transcriptomes against the *A. thaliana* mitHPPK/DHPS protein sequence. Sequences were aligned using Clustal Omega (Sievers et al., 2011) and rendered using the ESPRIPT 3.0 server (https://espript.ibcp.fr) (Robert and Gouet, 2014). Logos of selected sequences within catalytically relevant regions across 40 microbe and plant species (details in **Supplemental Tables 1**-**2**) were generated using the WebLogo server (Crooks et al., 2004).

### Expression and purification of cytHPPK/DHPS

*Escherichia coli* (Shuffle Express; NEB) containing the plasmid pREP4 (Qiagen) was transformed with an N-terminally His-tagged fusion protein of cytHPPK/DHPS (At1g69190) in the vector pQE30 (Qiagen). Overnight cultures grown at 30°C in LB broth supplemented with 35 µg/mL kanamycin and 100 µg/mL ampicillin were sub-cultured to an OD_600_ of 0.6 - 0.7 and induced overnight at 16°C with 100 mM isopropyl β-D-1-thiogalactopyranoside. The bacterial culture was centrifuged at 4,000 × *g* for 15 min at 4°C (Beckman) and the cell pellet was resuspended in 30-40 mL of lysis buffer (50 mM HEPES pH 8.0, 500 mM sodium chloride, 10 mM imidazole, 10 mM β-mercaptoethanol and 5% glycerol and lysozyme to final 1 mg/mL). After 30 min incubation on ice, the cell suspension was sonicated on ice for 6 min (40% amplitude, 3 s impulse, 9 s relapse) and centrifuged twice at 11,000 x *g* for 20 min at 4°C (Beckman) to pellet cell debris. The cell extract was clarified by filtering through 0.4 µm and 0.22 µm filters prior to loading onto ~4 mL of Ni-NTA resin (Profinity™ IMAC resin, Bio-Rad). To allow the His-tag protein to bind the resin it was incubated overnight at 4°C on a rolling shaker. To remove contaminant proteins, the resin was loaded into polypropylene gravity columns (Bio-Rad) and washed sequentially with 50 mL of ice-cold lysis buffer and 50 mL of washing buffer containing 50 mM HEPES pH 8.0, 500 mM sodium chloride, 30 mM imidazole, 10 mM β-mercaptoethanol and 5% glycerol. CytHPPK/DHPS was eluted from the columns using 10 mL of ice-cold elution buffer containing 50 mM HEPES pH 8.0, 500 mM sodium chloride, 250 mM imidazole, 10 mM β-mercaptoethanol and 5% glycerol. The eluate was centrifuged at 11,000 x *g* at 4°C to remove aggregated protein and the supernatant was concentrated by centrifugation on Amicon Ultra filters (MWCO 10 kDa) to a final volume of 0.2-0.3 mL. To remove excessive imidazole, the protein sample was diluted with ~30 mL of buffer containing 50 mM HEPES pH 8.0, 500 mM sodium chloride, 10 mM β-mercaptoethanol and 5% glycerol and concentrated again to a final volume of 0.6-0.8 mL. To remove aggregates and the remaining traces of imidazole, the protein sample was purified on an S200 size exclusion column (HiLoad® 16/600 Superdex® 200 pg) using 50 mM HEPES pH 8.0, 500 mM sodium chloride, 10 mM β-mercaptoethanol and 5% glycerol. The cytHPPK/DHPS peak was collected and concentrated by centrifugation on Amicon Ultra filters (MWCO 10 kDa) to a final volume of 0.2-0.3 mL and kept on ice before reductive methylation was carried out. A similar expression and purification approach was attempted for mitHPPK/DHPS (without its 68-residue N-terminal mitochondrial transit peptide), but we were unable to obtain soluble protein.

### Reductive methylation of cytHPPK/DHPS

To improve crystallisation of cytHPPK/DHPS, the surface lysine residues were reductively methylated according to the protocol described by Tan *et al*. (Tan et al., 2014). Purified cytHPPK/DHPS diluted to 16 mg/mL in 50 mM HEPES pH 8.0, 500 mM sodium chloride, 10 mM β-mercaptoethanol and 5% glycerol was mixed gently with 20 µL of 1 M dimethylamine-borane complex (Sigma) per 1 mL of protein solution. Immediately after addition of dimethylamine-borane complex, 40 µL of 1 M formaldehyde per 1 mL of protein solution was added to the reaction mixture and mixed gently. The solution was incubated at 4°C for two hours and the procedure was repeated. After two hours of incubation, an additional 10 µL of 1 M dimethylamine-borane complex per 1 mL of protein solution was added to the reaction mixture which was incubated at 4°C for 12-14 hours. To quench the reaction, 80 µL of 1 M glycine and 6 µL of 1 M dithiothreitol were added to the reaction mixture, and the solution was left on ice for two hours. To remove unreacted dimethylamine-borane complex, formaldehyde and high molecular weight protein aggregates, the reaction mixture was purified by size-exclusion chromatography on an S200 size exclusion column (HiLoad® 16/600 Superdex® 200 pg) and eluted in 20 mM HEPES pH 8.0, 250 mM sodium chloride, 2 mM dithiothreitol. The cytHPPK/DHPS peak was collected and concentrated by centrifugation in an Amicon Ultra filter (MWCO 10 kDa). To remove excess salt, the sample was diluted with 20 mM HEPES pH 8.0, 125 mM sodium chloride, 2 mM dithiothreitol and concentrated to ~10–20 mg/mL for use in crystallisation trials.

### Crystallisation and structure determination of *A. thaliana* cytHPPK/DHPS

Crystals of cytHPPK/DHPS were grown at 20°C using the sitting drop vapour-diffusion method by mixing reservoir buffer (30-40% PEG 3350 and 0.2 M sodium fluoride) with protein solution (13–20 mg/mL) in a 1:3 or 1:4 ratio. To obtain crystals of protein-herbicide complex, a single droplet containing several cytHPPK/DHPS crystals in reservoir buffer was soaked for 1 week with asulam (by dusting powdered herbicide on to a droplet). Prior to screening, crystals were cryo-protected in 25% glycerol, 30-40% PEG 3350 and 0.2 M sodium fluoride, and flash-cooled in liquid nitrogen. X-ray data were collected using the MX2 beamline at the Australian Synchrotron, part of Australia’s Nuclear Science and Technology Organisation (ANSTO) (Aragao et al., 2018). The X-ray data were indexed using iMosflm (Battye et al., 2011) and space group was assigned using Pointless (CCP4 package) (Winn et al., 2011). Unmerged data were further processed using the STARANISO server (Tickle, 2018) to scale, merge and correct for anisotropy. The cytHPPK/DHPS structure was determined by molecular replacement using PHASER (from within the PHENIX package) (Adams et al., 2010) and an ensemble model generated in Chainsaw (CCP4 package) (Winn et al., 2011) using *S. cerevisiae* HPPK/DHPS (PDB ID: 2bmb). The molecular replacement model was subsequently rebuilt using PHENIX.AUTOBUILD (Terwilliger et al., 2008) and refined in Refmac (CCP4 package)(Winn et al., 2011). Coordinates and structure factors for cytHPPK/DHPS were deposited into the Protein Data Bank (PDB) under accession code 7mpy.

### Chemical synthesis of pterin-mimic inhibitors

Compounds **8**-**12** were synthesised as previously described (Dennis et al., 2014; Yun et al., 2014; Dennis et al., 2016; Dennis et al., 2018)and purified by C18 chromatography (0-100% acetonitrile/0.1% trifluoracetic acid in water) (9-19%).

### Minimum inhibitory concentration (MIC) assay

MICs were performed in triplicate using the broth microdilution method (Clinical and Laboratory Standards Institute) (CLSI, 2021) in a 96-well plate with serial dilutions of sulfonamides (AK Scientific), pterin-mimicking inhibitors (synthesis described above), glyphosate (Sigma-Aldrich) and oryzalin (Sigma-Aldrich) in 100 μL of cation-adjusted Mueller–Hinton broth. To account for the dimethyl sulfoxide used to dissolve HPPK/DHPS inhibitors, an equivalent quantity was added to the Mueller–Hinton broth to a final 0.6-2% concentration. Wells were inoculated with 100 μL of ~10^5^ cells of *E. coli* or *B. subtilis* grown to an optical density (OD_600_) of ~0.5, with the final inhibitor concentration ranging from 0.04 – 2 mM. MICs were determined by visualizing no growth wells after incubation of the cells for 20 hours at 37 °C in a shaker incubator.

### Sulfonamide treatments on soil-grown *A. thaliana*

Approximately 60 *A. thaliana* Col-0 seeds were sown in pots (63 × 63 × 59 mm) of Seedling Substrate Plus+ soil (Bord na Móna Horticulture Ltd, Newbridge, Ireland) consisting of Irish peat. Soil was pre-wet before sowing to saturation and then watered accordingly throughout the experiment to maintain adequate moist conditions. No fertiliser was added to the soil. Seeds were cold-treated for 3 days in the dark at 4°C to synchronise germination and then grown in a chamber at 23°C, with 55% RH and in a 16 h light/8 h dark photoperiod. Sulfonamides (AK Scientific), glyphosate (Sigma Aldrich) and oryzalin (Sigma Aldrich) were initially dissolved in DMSO at 20 mg/mL and further diluted in water prior to treatments. The surfactant Brushwet (SST Dandenong, Australia) was added to a final concentration of 0.2 mL/L. The active ingredients glyphosate and oryzalin of commercial herbicides were used as positive controls. Each compound was tested on seeds (pre-emergence) or seedlings (post emergence) by spraying each pot with 0.5 mL of 0.125, 0.25, 0.5, 1, 2, or 4 mM solutions directly onto the plants, which are equivalent to ~27-1,562 g/ha a.i. The soil density in each pot was 0.29 g/cm^3^, based on dry soil, and the total volume of herbicide solution per gram of soil was 10 µL/g for pre-emergence and 20 µL/g for post emergence, because a single application was used for pre-emergence (0.5 mL), whereas two applications of 0.5 mL were used post emergence. The day the trays were moved into their first long day was considered as day 0 for which the single pre-emergence treatments were given. Post-emergence treatments were carried out twice post germination, at days 4 and 8, and seedlings were grown for 15 days before photographs were taken. The herbicidal effect of each compound was assessed visually.

## Acknowledgements

This research was undertaken in part using the MX2 beamline at the Australian Synchrotron, part of ANSTO, and made use of the Australian Cancer Research Foundation (ACRF) detector. K.V.S. was supported by the Australian Research Training Program scholarship. This work and G.V. were supported by Australian Research Council grant DP190101048 to J.S.M., K.A.S. and J.H. who was also supported by an ARC Discovery Early Career Researcher Award (grant no. DE180101445).

## Author contribution statement

J.S.M., G.V., and K.V.S. conceived the study. K.V.S. performed recombinant protein purification and crystallisation, and G.V. determined the crystal structure with J.H. and C.S.B. providing input. G.V., J.H., J.S.M. and K.V.S. performed plant and microbial sensitivity testing. K.J.B. and K.A.S. performed chemical syntheses of mercaptoguanine inhibitors. M.F.F. assembled transcriptomes of weed species and determined their HPPK/DHPS sequences. G.V. and J.S.M. wrote the manuscript with input from all authors.

## Supplemental Information

**Supplemental Figure 1.**
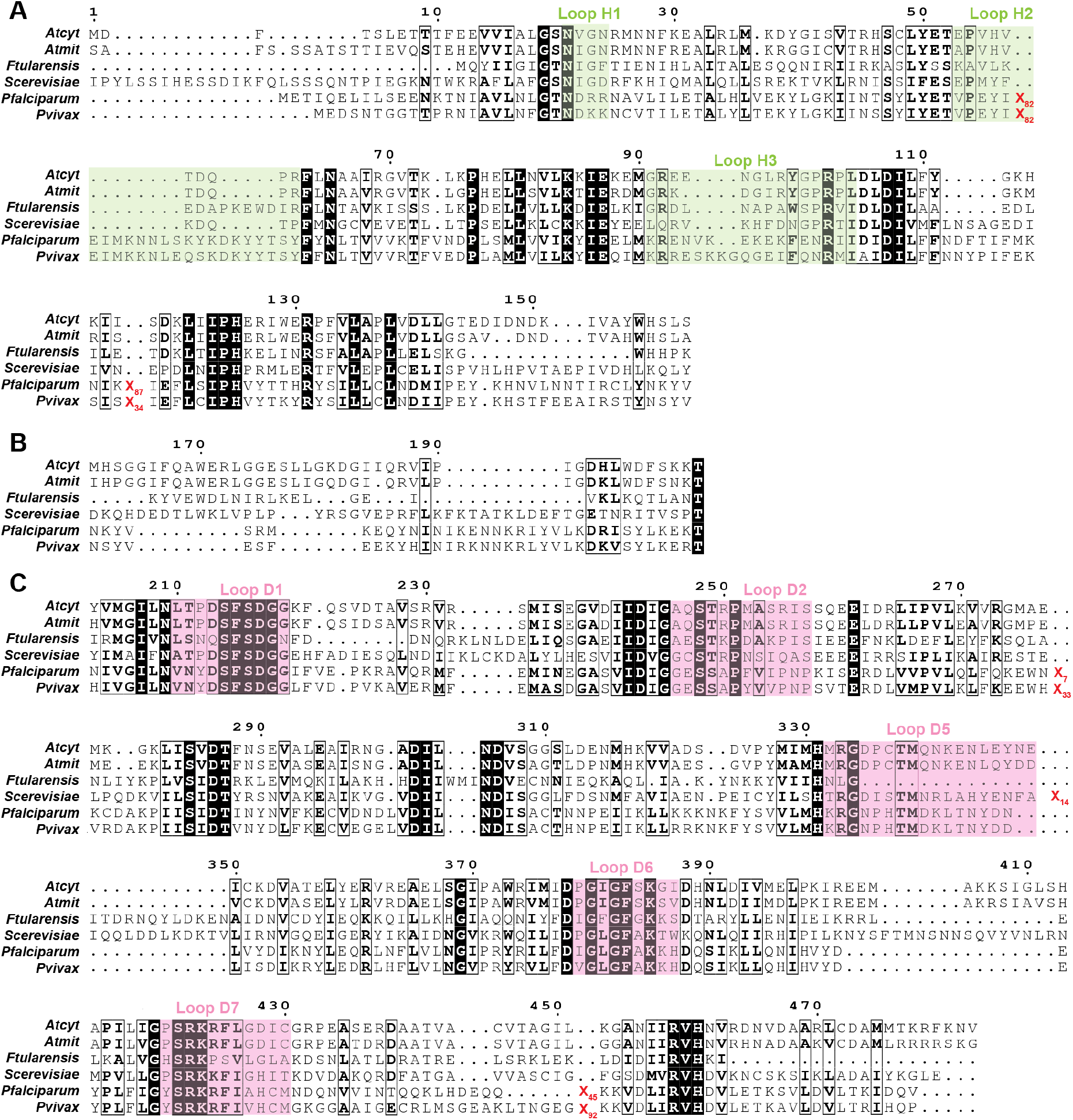
Bifunctional HPPK/DHPS from *S. cerevisiae, F. tularensis, P. falciparum*, and *P. vivax* contain insertions of varying lengths relative to the *A. thaliana* enzyme. Alignments show sequences of **(A)** HPPK, **(B)** the linker region, and **(C)** DHPS. Long inserts within the HPPK and DHPS domains in other species are represented by X_n_ (n = number of residues). These inserts occur outside of the core active site residues involved in catalysis. Conserved and similar residues are shown in white and bold font, respectively. Catalytic loops in HPPK and DHPS are highlighted in green and pink, respectively.

**Supplemental Figure 2.**
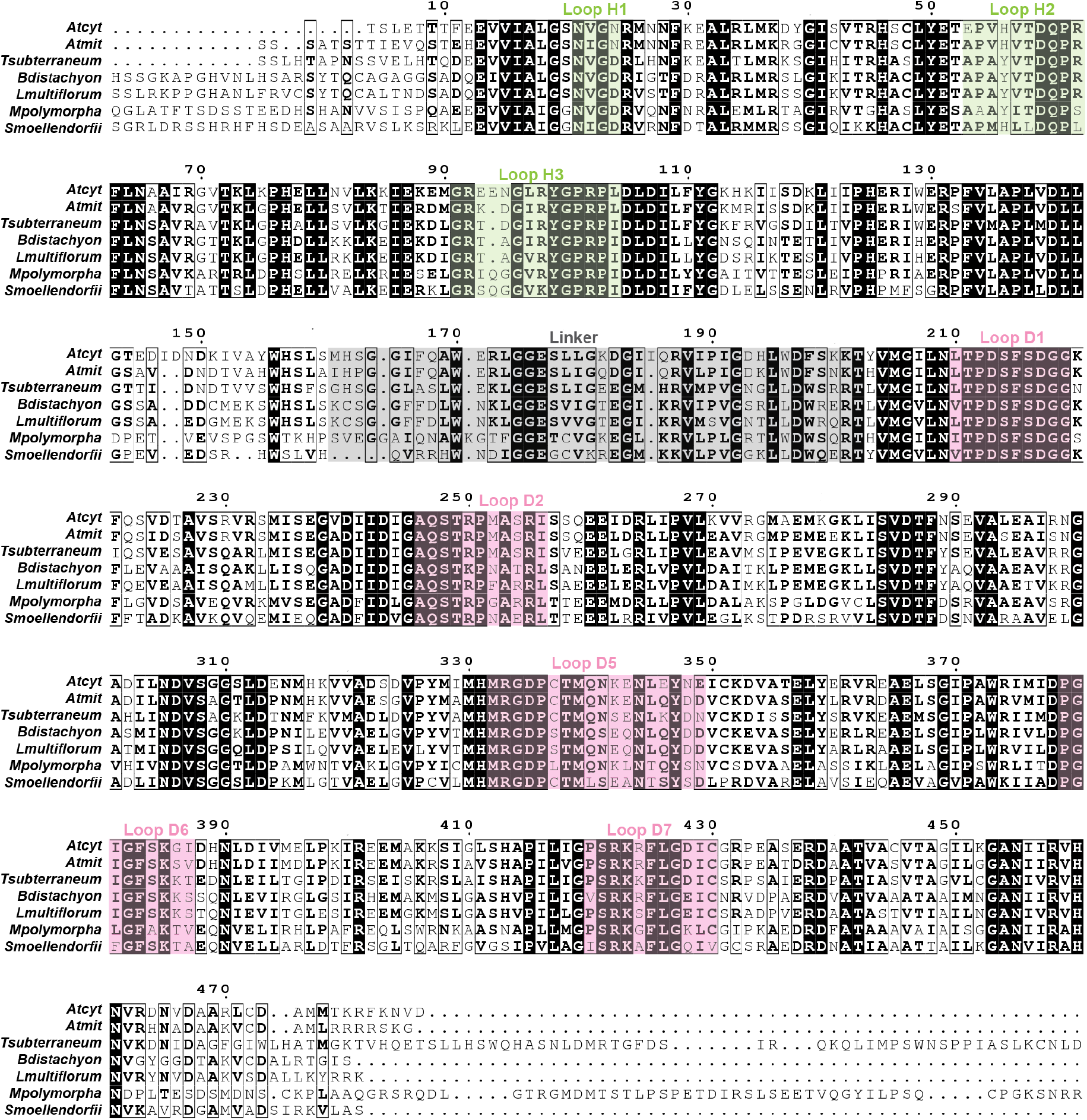
Active site regions are conserved in bifunctional HPPK/DHPS across the plant kingdom. Sequences are shown for *A. thaliana* HPPK/DHPS homologs, *Trifolium subterraneum, Brachypodium distachyon, Lolium multiflorum, Marchantia polymorpha*and *Selaginella moellendorfii*. Conserved and similar residues are shown in white and bold font, respectively. Catalytic loop regions in HPPK and DHPS are highlighted green and pink, respectively. The linker region is highlighted in grey.

**Supplemental Figure 3.**
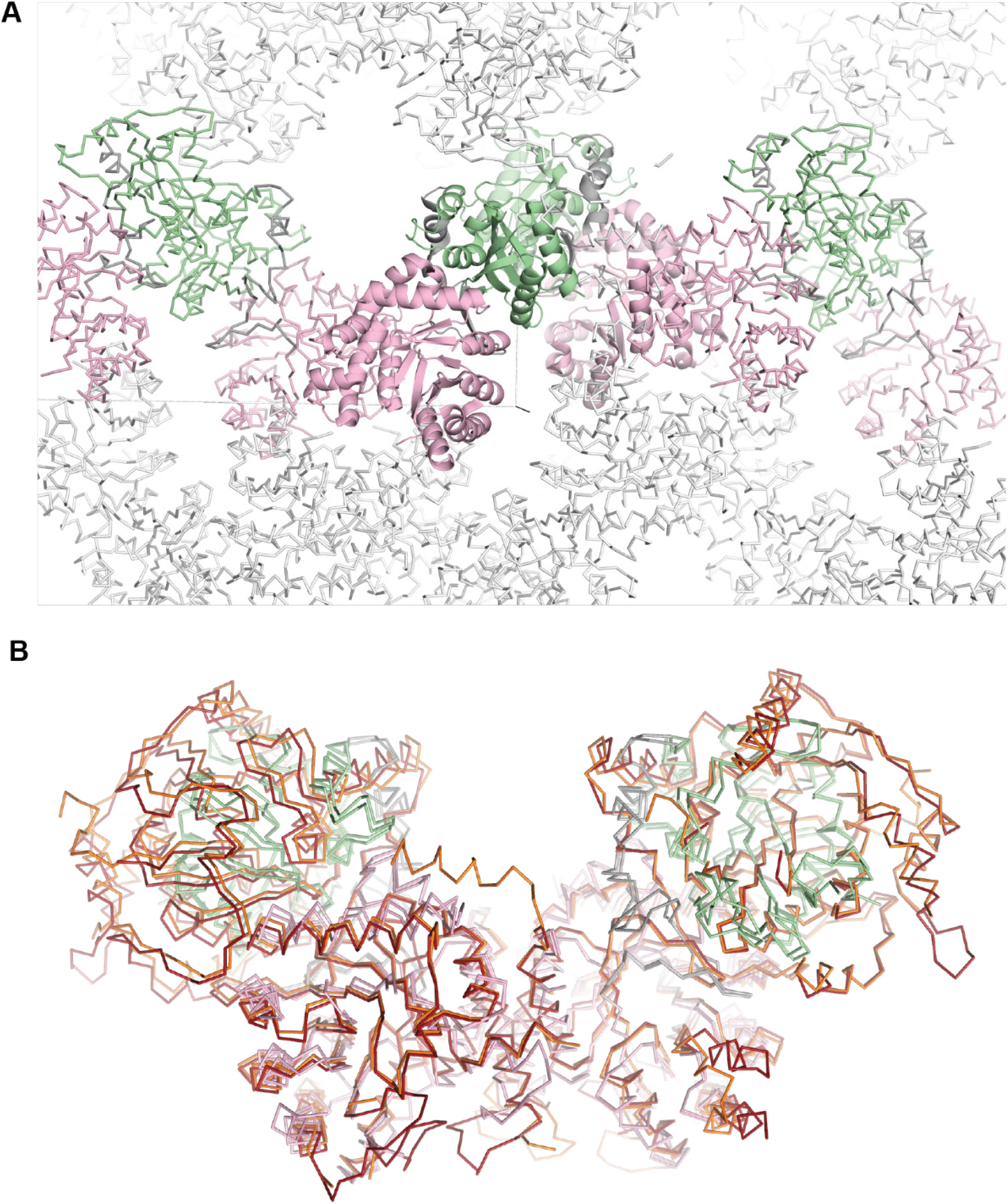
cytHPPK/DHPS dimerises similarly to HPPK/DHPS homologs. **(A)** The cytHPPK/DHPS crystal lattice showing the association between chains A:B within the asymmetric unit (shown as cartoon), and their dimeric association with symmetry mates A’ and B’ (shown as ribbons). **(B)** Superposition of cytHPPK/DHPS dimers A:A’ and B:B’ with *P. vivax* HPPK/DHPS (PDB ID: 5z79; red) and *P. falciparum* HPPK/DHPS (PDB ID: 6jwq; orange) showing similar association between dimers.

**Supplemental Figure 4.**
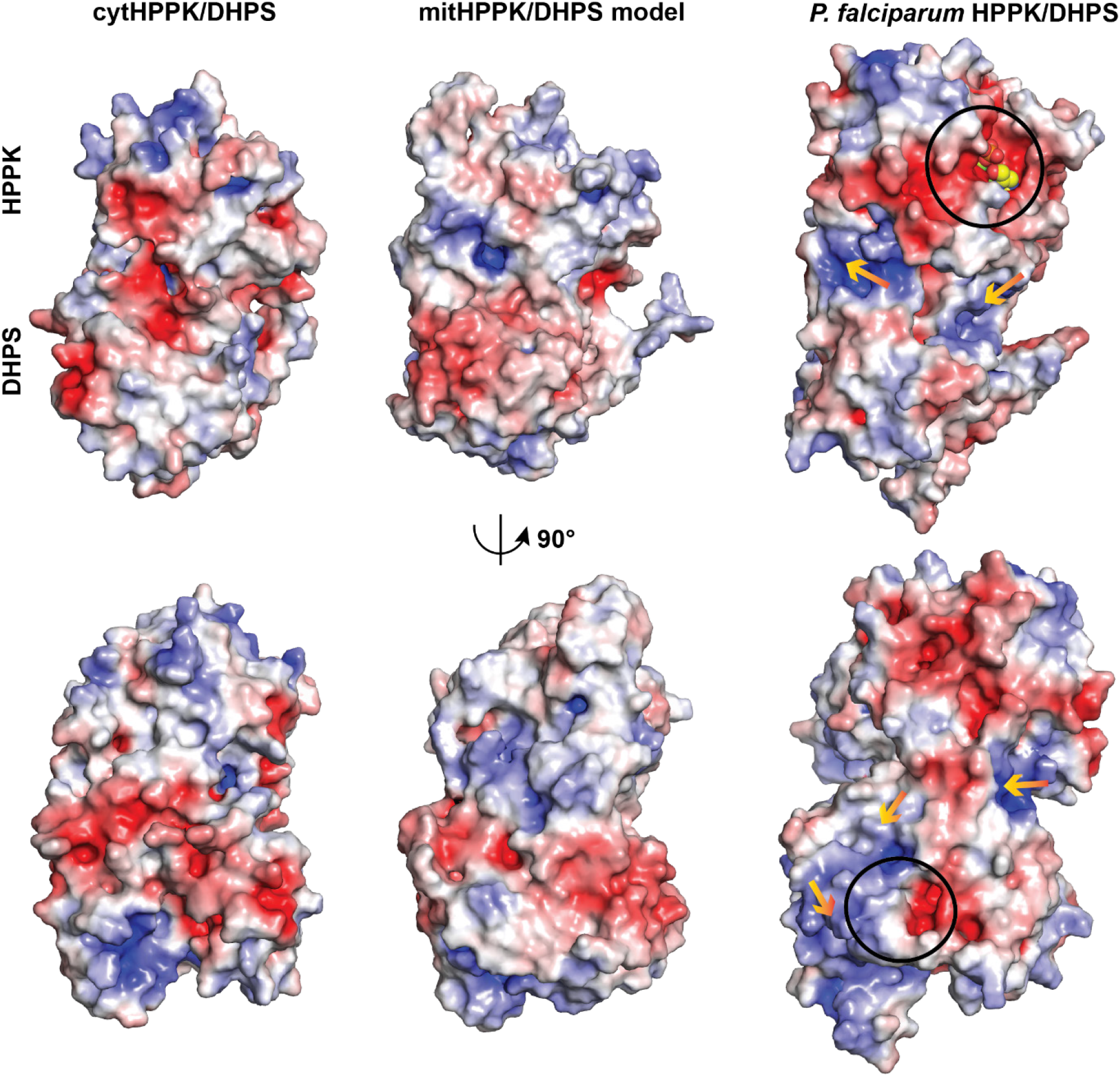
*A. thaliana* HPPK/DHPS has no obvious positively charged channel between active sites. The electrostatic potential surface map for *A. thaliana* cytHPPK/DHPS (chain B) structure was superposed with a model of mitHPPK/DHPS (generated using cytHPPK/DHPS as a reference model by iTASSER (Yang et al., 2015a)), and *P. falciparum* HPPK/DHPS (PDB ID: 6jwr). HPPK and DHPS active sites in *Pf*HPPK/DHPS are circled, with bound substrate shown as spheres. Arrows indicate a proposed substrate channelling path from the HPPK active site to the DHPS active site in *Pf*HPPK/DHPS (Chitnumsub et al, 2020). Electronegative and electropositive surfaces are rendered red and blue, respectively within the range of −5.0 and 5.0. Images were prepared using Pymol v2.5.2 (Schrödinger, 2015).

**Supplemental Figure 5.**
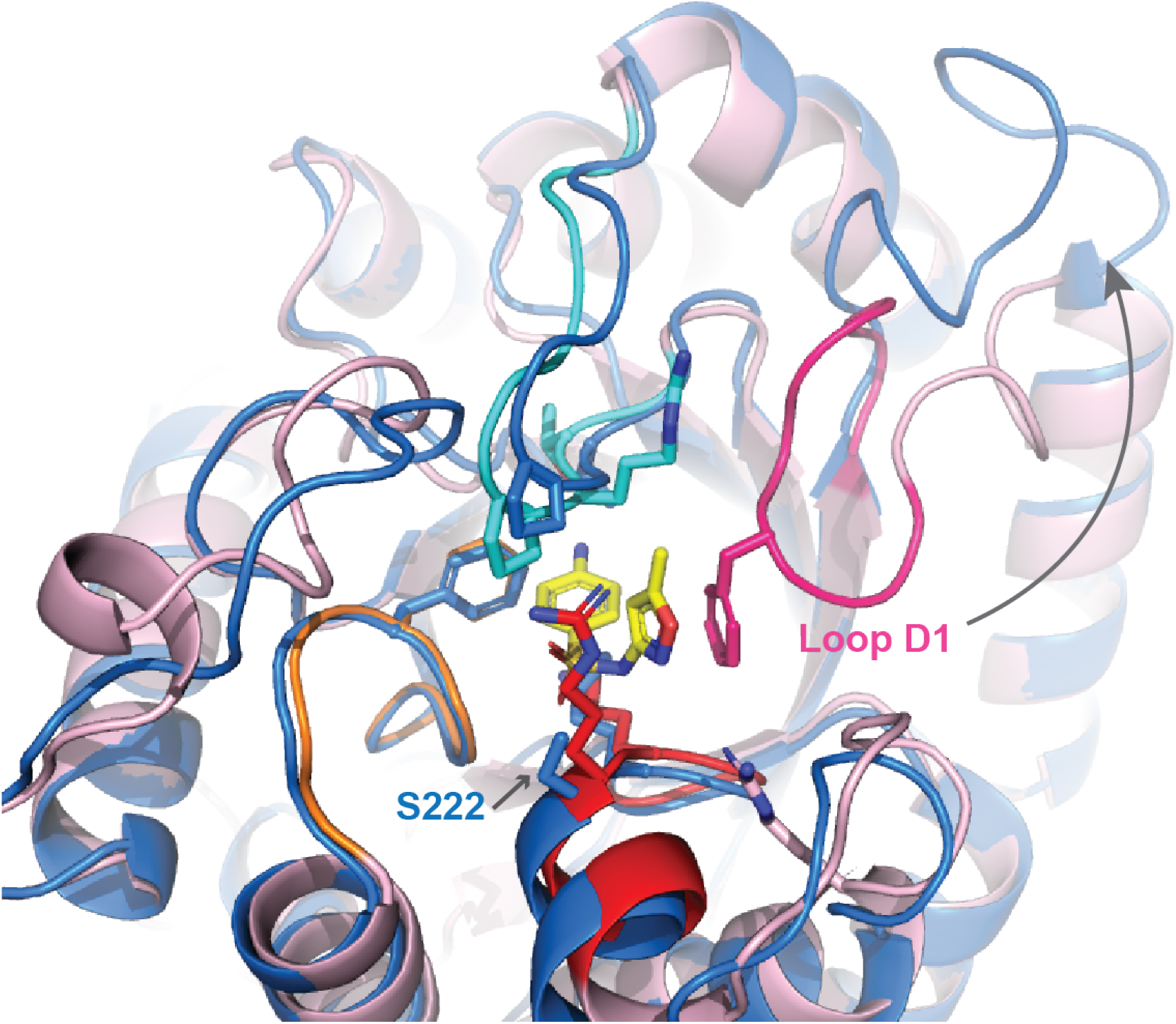
DHPS catalytic loop D1 adopts conformations away from the active site in some structures. Superposition of *A. thaliana* sulfamethoxazole-bound mitDHPS model with *E. coli* DHPS (PDB ID: 1ajz) shows loop D1 in a conformation away from the *p*-ABA binding pocket. Ligand atoms are shown in yellow (carbon), blue (nitrogen), and red (oxygen), respectively.

### Pterin-based compound treatments on soil-grown *A. thaliana*

Approximately 30 *A. thaliana* Col-0 seeds were sown in pots (63 × 63 × 59 mm) of Seedling Substrate Plus+ soil (Bord na Móna Horticulture Ltd, Newbridge, Ireland) consisting of Irish peat. Soil was pre-wetted to saturation before sowing and then watered accordingly throughout the experiment to maintain adequate moist conditions. No fertiliser was added to the soil. Seeds were cold-treated for 3 days in the dark at 4°C to synchronise germination and then grown in a chamber at 22°C, with 60% RH and in a 16 h light/8 h dark photoperiod. Herbicides and test compounds were initially dissolved in DMSO at 20 mg/mL and further diluted in water prior to treatments. The surfactant Brushwet (SST Dandenong, Australia) was added to a final concentration of 0.02%. The active ingredients glyphosate and oryzalin of commercial herbicides were used as positive controls.

Herbicidal activity was assessed in triplicate at each concentration of pterin mimic inhibitors. The negative control contained 2% DMSO and 0.02% Brushwet. Each compound was tested on seeds (pre-emergence) or seedlings (post emergence) by pipetting 500 µL of 0, 25, 50, 100, 200 or 400 mg/L solutions directly onto the plants; these quantities and concentrations which are equivalent to ~35, 70, 140, 280 and 560 g/ha a.i. The soil density in each pot was 0.29 g/cm^3^, based on dry soil, and the total volume of herbicide solution per gram of soil was 10 µL/g for pre-emergence and 20 µL/g for post emergence, because a single application was used for pre-emergence (0.5 mL), whereas two applications of 0.5 mL were used post emergence. The day the trays were moved into their first long day was considered as day 0 for which the single pre-emergence treatments were given. Post-emergence treatments were carried out twice post germination, at days 3 and 6, and seedlings were grown for 16 days before photographs were taken. The herbicidal effect of each compound was assessed visually.

**Supplemental Figure 6.**
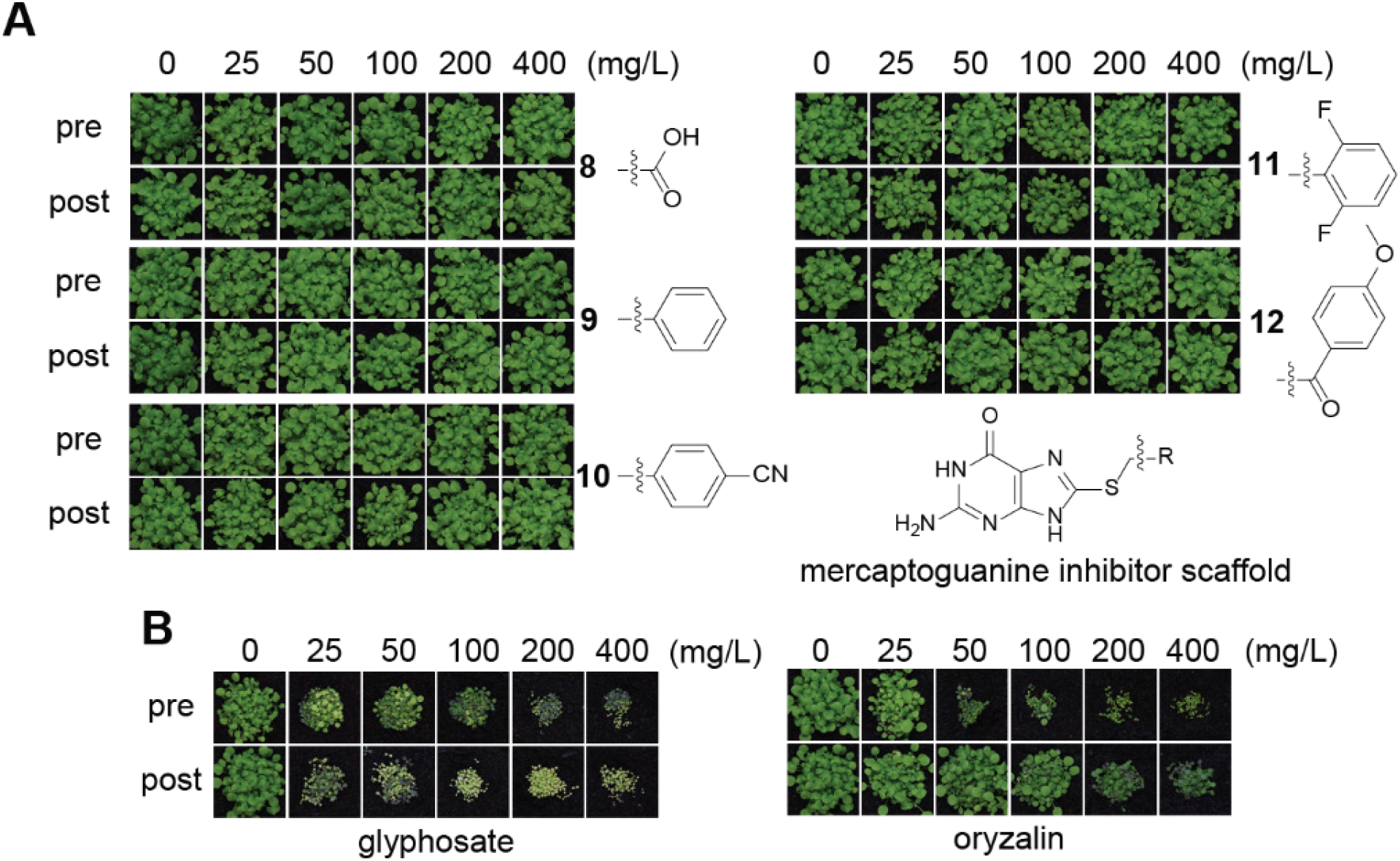
Mercaptoguanine-based HPPK/DHPS inhibitors are not herbicidal. Pre- and post-emergence herbicidal activities against *A. thaliana* are shown for **(A)** mercaptoguanine-based inhibitors 8-12 (named in **Supplemental Table 3**), and **(B)** herbicide controls glyphosate and oryzalin.

**Supplemental Figure 7.**
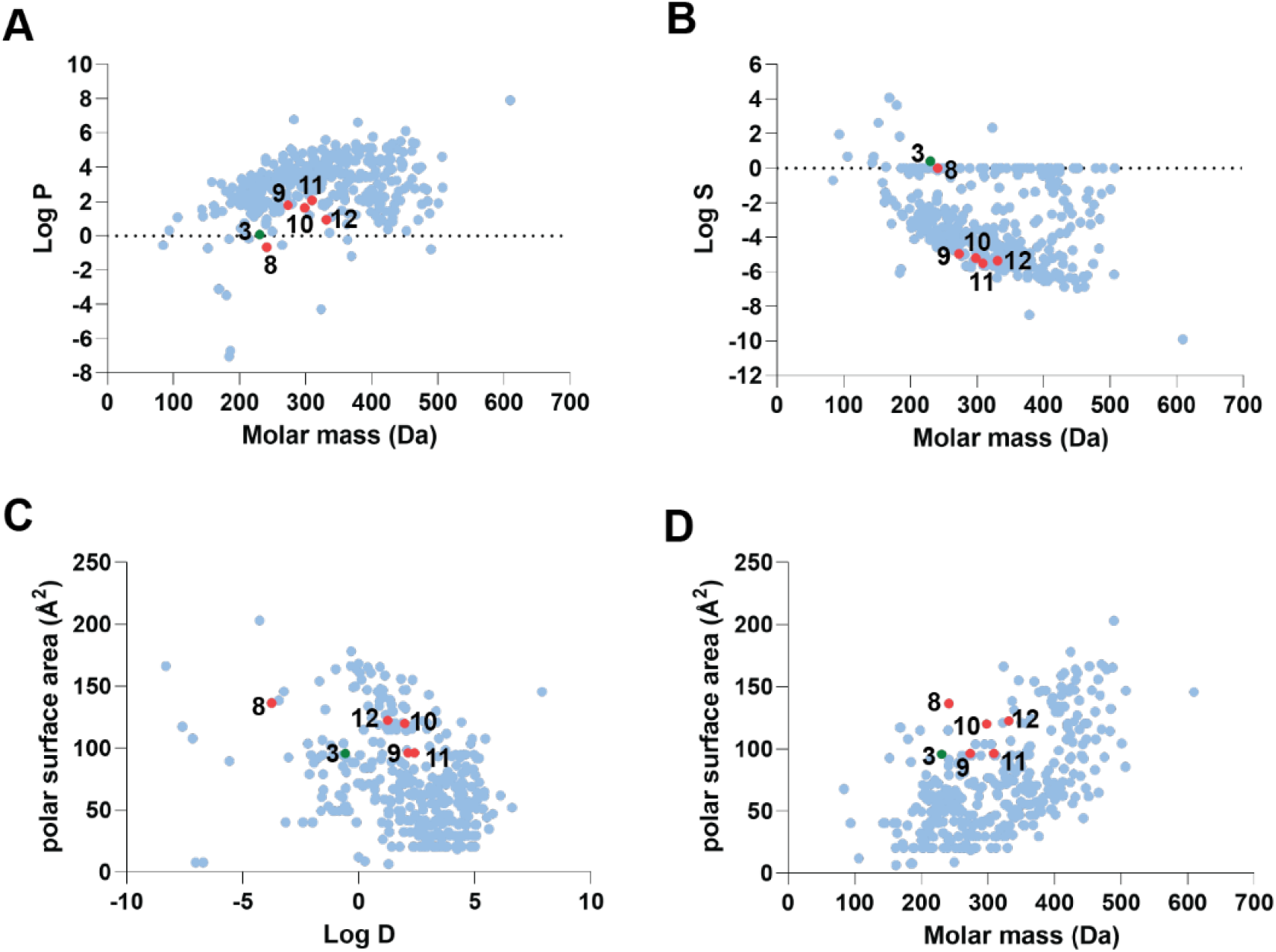
Physicochemical properties of mercaptoguanine-based HPPK/DHPS inhibitors. Physicochemical data for mercaptoguanine-based inhibitors (red), asulam (green) and 359 commercial herbicides (blue) using an interactive database by Sukhoverkov et al. (Sukhoverkov et al., 2021) with graphs comparing **(A)** molar mass *vs* lipophilicity (Log *P*), **(B)** molar mass *vs* aqueous solubility (Log *S*), **(C)** distribution coefficient (Log *D*) *vs* polar surface area (Å^2^), **(D)** molar mass *vs* polar surface area (Å^2^).

**Supplemental Table 1.**
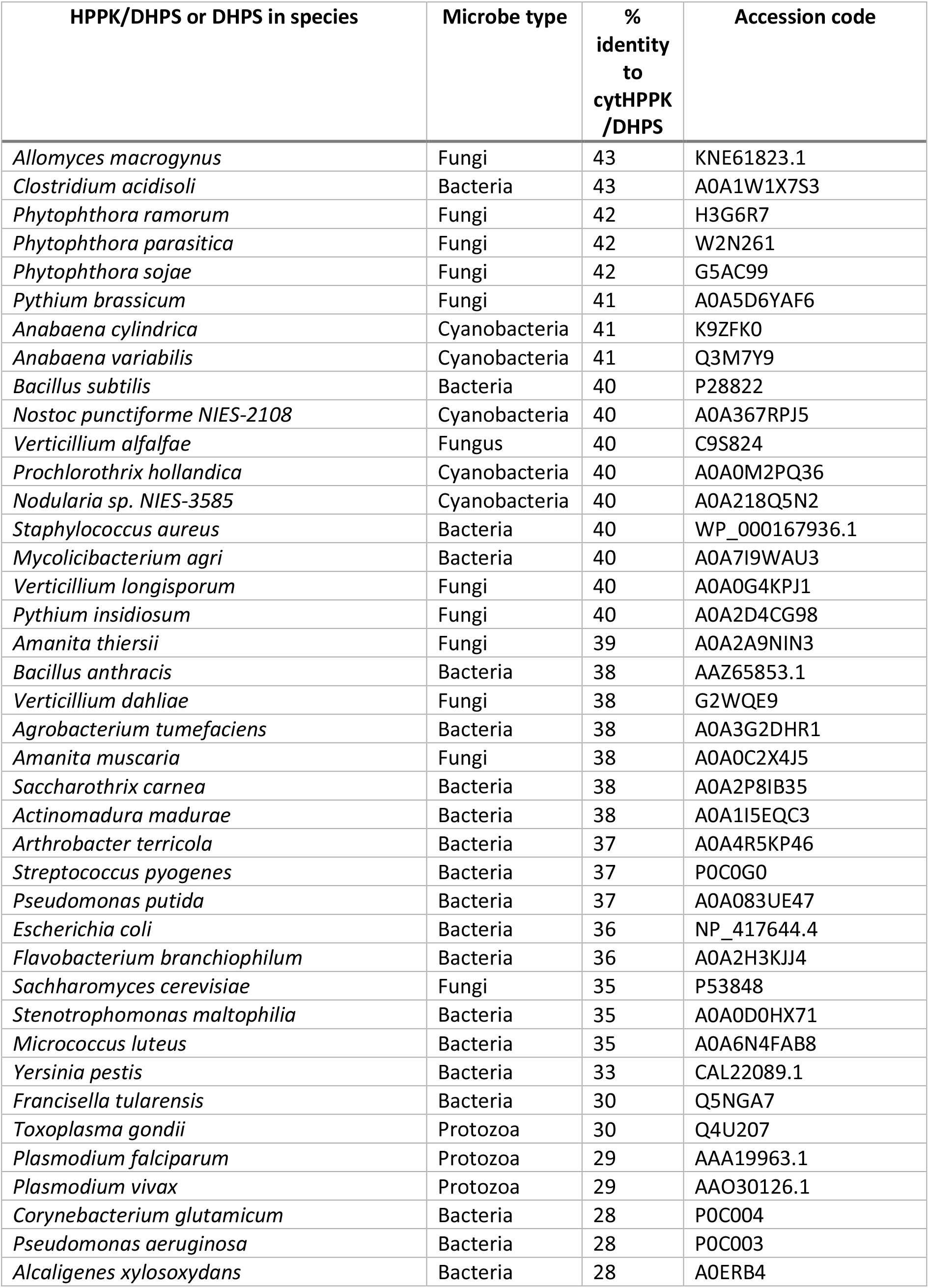
Bifunctional HPPK-DHPS and monofunctional DHPS sequences from microbes.

**Supplemental Table 2.** Bifunctional HPPK/DHPS sequences from plants.

| HPPK/DHPS in species | Clade | % identity to cytHPPK/DHPS | Accession code |
| --- | --- | --- | --- |
| *Arabidopsis thaliana* (cytosolic) | Eudicot | 100 | Q1ENB6 |
| *Arabidopsis arenosa* | Eudicot | 80 | CAE6168687.1 |
| *Arabidopsis thaliana* (mitochondrial) | Eudicot | 80 | Q9SZV3 |
| *Arabidopsis suecica* | Eudicot | 80 | KAG7545658.1 |
| *Arabidopsis lyrata subsp lyrata* | Eudicot | 80 | XP_002867374.1 |
| *Arabis alpina* | Eudicot | 79 | KFK29559.1 |
| *Brassica rapa* | Eudicot | 77 | XP_033134457.1 |
| *Populus trichocarpa* | Eudicot | 69 | XP_024462420.1 |
| *Vitis vinifera* | Eudicot | 68 | RVW42188.1 |
| *Lupinus angustifolius* | Monocot | 68 | XP_019428656.1 |
| *Gossypium barbadense* | Eudicot | 67 | PPD74637.1 |
| *Eucalyptus grandis* | Eudicot | 67 | XP_010062483.2 |
| *Malus domestica* | Eudicot | 67 | XP_008383585.1 |
| *Pisum sativum* | Eudicot | 66 | O04862 |
| *Glycine max* | Eudicot | 66 | XP_003518782.1 |
| *Ricinus communis* | Eudicot | 66 | B9S4E5 |
| *Medicago truncatula* | Eudicot | 66 | XP_003614113.1 |
| *Ipomoea triloba* | Eudicot | 65 | XP_031097973.1 |
| *Trifolium subterraneum* | Eudicot | 65 | GAU13272.1 |
| *Spinacia oleracea* | Eudicot | 65 | XP_021845571.1 |
| *Solanum lycopersicum* | Eudicot | 65 | NP_001317113.1 |
| *Solanum tuberosum* | Eudicot | 65 | XP_006348669.1 |
| *Cocos nucifera* | Monocot | 64 | KAG1347762.1 |
| *Chenopodium quinoa* | Eudicot | 63 | XP_021756582.1 |
| *Digitaria exilis* | Monocot | 62 | KAF8724376.1 |
| *Sorghum bicolor* | Monocot | 62 | XP_021313165.1 |
| *Triticum aestivum* | Monocot | 61 | ABP01351.1 |
| *Setaria viridis* | Monocot | 61 | XP_034568867.1 |
| *Triticum turgidum subsp durum* | Monocot | 61 | VAH43760.1 |
| *Cyperus rotundus* | Monocot | 61 | Assembled from RNASeq |
| *Lolium multiflorum* | Monocot | 61 | Assembled from RNASeq |
| *Oryza sativa* | Monocot | 60 | Q7X7X0 |
| *Brachypodium distachyon* | Monocot | 60 | I1GS77 |
| *Echinochloa crus-galli* | Monocot | 60 | Assembled from RNASeq |
| *Zea mays* | Monocot | 60 | NP_001146490.1 |
| *Ceratodon purpureus* | Moss | 53 | KAG0575880.1 |
| *Physcomitrium patens* | Moss | 53 | XP_024367173.1 |
| *Marchantia paleacea* | Marchantiophyta | 52 | KAG6554302.1 |
| *Marchantia polymorpha* | Marchantiophyta | 50 | PTQ44222.1 |
| *Selaginella moellendorffii* | Club-moss | 48 | XP_002963403.2 |

**Supplemental Table 3.**
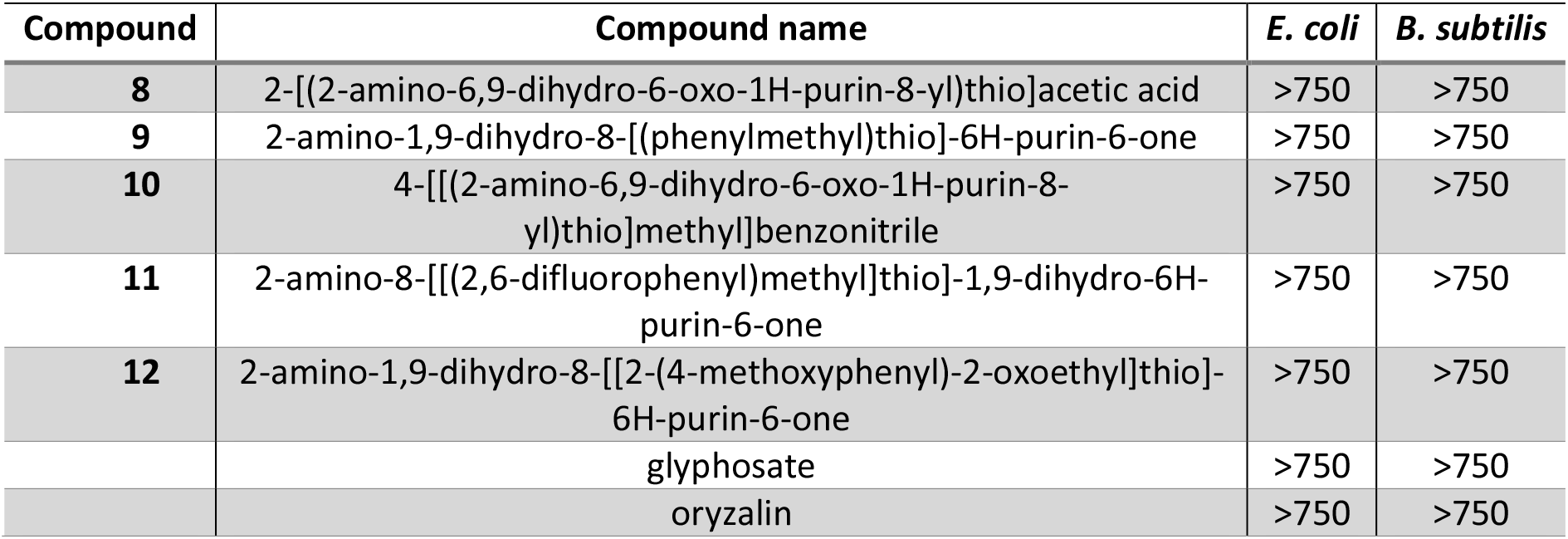
Minimum inhibitory concentration (μM) of pterin-mimicking HPPK/DHPS inhibitors.

## References

Achari, A., Somers, D.O., Champness, J.N., Bryant, P.K., Rosemond, J., and Stammers, D.K. (1997). Crystal structure of the anti-bacterial sulfonamide drug target dihydropteroate synthase. Nat. Struct. Biol. 4, 490–497.

Adams, P.D., Afonine, P.V., Bunkóczi, G., Chen, V.B., Davis, I.W., Echols, N., Headd, J.J., Hung, L.W., Kapral, G.J., and Grosse-Kunstleve, R.W. (2010). PHENIX: a comprehensive Python-based system for macromolecular structure solution. Acta Crystallographica Section D: Biological Crystallography 66, 213–221.

Anderson, K.S. (2017). Understanding the molecular mechanism of substrate channeling and domain communication in protozoal bifunctional TS-DHFR. Protein Eng. Des. Sel. 30, 253–261.

Aragao, D., Aishima, J., Cherukuvada, H., Clarken, R., Clift, M., Cowieson, N.P., Ericsson, D.J., Gee, C.L., Macedo, S., Mudie, N., Panjikar, S., Price, J.R., Riboldi-Tunnicliffe, A., Rostan, R., Williamson, R., and Caradoc-Davies, T.T. (2018). MX2: a high-flux undulator microfocus beamline serving both the chemical and macromolecular crystallography communities at the Australian Synchrotron. J. Synchrotron Radiat. 25, 885–891.

Arnaud, G., Colin, L., and Jeremy, P.D. (2006). Crystal structure of the bifunctional dihydroneopterin aldolase/6-hydroxymethyl-7,8-dihydropterin pyrophosphokinase from Streptococcus pneumoniae. J. Mol. Biol. 360, 644–653.

Babaoglu, K., Qi, J., Lee, R.E., and White, S.W. (2004). Crystal structure of 7,8-dihydropteroate synthase from Bacillus anthracis: mechanism and novel inhibitor design. Structure 12, 1705–1717.

Battye, T.G.G., Kontogiannis, L., Johnson, O., Powell, H.R., and Leslie, A.G.W. (2011). iMOSFLM: a new graphical interface for diffraction-image processing with MOSFLM. Acta Crystallogr. D 67, 271–281.

Blaszczyk, J., Shi, G., Yan, H., and Ji, X. (2000). Catalytic center assembly of HPPK as revealed by the crystal structure of a ternary complex at 1.25 Å resolution. Structure 8, 1049–1058.

Bourne, C.R. (2014). Utility of the biosynthetic folate pathway for targets in antimicrobial discovery. Antibiotics (Basel) 3, 1–28.

Brain, R.A., Ramirez, A.J., Fulton, B.A., Chambliss, C.K., and Brooks, B.W. (2008). Herbicidal effects of sulfamethoxazole in Lemna gibba: using p-aminobenzoic acid as a biomarker of effect. Environ. Sci. Technol. 42, 8965–8970.

Chakraborty, S., Gruber, T., Barry, C.E., Boshoff, H.I., and Rhee, K.Y. (2013). Para-aminosalicylic acid acts as an alternative substrate of folate metabolism in Mycobacterium tuberculosis. Science 339, 88–91.

Chen, S., McElroy, J.S., Dane, F., and Goertzen, L.R. (2016). Transcriptome assembly and comparison of an allotetraploid weed species, annual bluegrass, with its two diploid progenitor species, Poa supina Schrad and Poa infirma Kunth. Plant Genome 9, 1–11.

Cheong, M.S., Seo, K.H., Chohra, H., Yoon, Y.E., Choe, H., Kantharaj, V., and Lee, Y.B. (2020). Influence of sulfonamide contamination derived from veterinary antibiotics on plant growth and development. Antibiotics (Basel) 9, 456.

Chhabra, S., Barlow, N., Dolezal, O., Hattarki, M.K., Newman, J., Peat, T.S., Graham, B., and Swarbrick, J.D. (2013). Exploring the chemical space around 8-mercaptoguanine as a route to new inhibitors of the folate biosynthesis enzyme HPPK. PLoS One 8, e59535.

Chitnumsub, P., Jaruwat, A., Talawanich, Y., Noytanom, K., Liwnaree, B., Poen, S., and Yuthavong, Y. (2020). The structure of Plasmodium falciparum hydroxymethyldihydropterin pyrophosphokinase-dihydropteroate synthase reveals the basis of sulfa resistance. FEBS J. 287, 3273–3297.

CLSI. (2021). Performance standards for antimicrobial susceptibility testing, 31st Edition M100-S31 (Wayne, Pennsylvania: Clinical and Laboratory Standards Institute).

Cossins, E.A., and Chen, L. (1997). Folates and one-carbon metabolism in plants and fungi. Phytochemistry 45, 437–452.

Crooks, G.E., Hon, G., Chandonia, J.-M., and Brenner, S.E. (2004). WebLogo: A sequence logo generator. Genome Res. 14, 1188–1190.

Czaban, A., Sharma, S., Byrne, S.L., Spannagl, M., Mayer, K.F., and Asp, T. (2015). Comparative transcriptome analysis within the Lolium/Festuca species complex reveals high sequence conservation. BMC Genomics 16, 1–13.

Dennis, M.L., Pitcher, N.P., Lee, M.D., DeBono, A.J., Wang, Z.C., Harjani, J.R., Rahmani, R., Cleary, B., Peat, T.S., Baell, J.B., and Swarbrick, J.D. (2016). Structural basis for the selective binding of inhibitors to 6-Hydroxymethyl-7,8-dihydropterin pyrophosphokinase from Staphylococcus aureus and Escherichia coli. J. Med. Chem. 59, 5248–5263.

Dennis, M.L., Chhabra, S., Wang, Z.C., Debono, A., Dolezal, O., Newman, J., Pitcher, N.P., Rahmani, R., Cleary, B., Barlow, N., Hattarki, M., Graham, B., Peat, T.S., Baell, J.B., and Swarbrick, J.D. (2014). Structure-based design and development of functionalized Mercaptoguanine derivatives as inhibitors of the folate biosynthesis pathway enzyme 6-hydroxymethyl-7,8-dihydropterin pyrophosphokinase from Staphylococcus aureus. J. Med. Chem. 57, 9612–9626.

Dennis, M.L., Lee, M.D., Harjani, J.R., Ahmed, M., DeBono, A.J., Pitcher, N.P., Wang, Z.C., Chhabra, S., Barlow, N., Rahmani, R., Cleary, B., Dolezal, O., Hattarki, M., Aurelio, L., Shonberg, J., Graham, B., Peat, T.S., Baell, J.B., and Swarbrick, J.D. (2018). 8-mercaptoguanine derivatives as inhibitors of dihydropteroate synthase. Chemistry 24, 1922–1930.

European Food Safety Authority, Arena, M., Auteri, D., Barmaz, S., Brancato, A., Brocca, D., Bura, L., Chiusolo, A., Court Marques, D., Crivellente, F., De Lentdecker, C., Egsmose, M., Fait, G., Ferreira, L., Goumenou, M., Greco, L., Ippolito, A., Istace, F., Jarrah, S., Kardassi, D., Leuschner, R., Lythgo, C., Magrans, J.O., Medina, P., Miron, I., Molnar, T., Nougadere, A., Padovani, L., Parra Morte, J.M., Pedersen, R., Reich, H., Sacchi, A., Santos, M., Serafimova, R., Sharp, R., Stanek, A., Streissl, F., Sturma, J., Szentes, C., Tarazona, J., Terron, A., Theobald, A., Vagenende, B., and Villamar-Bouza, L. (2018). Peer review of the pesticide risk assessment of the active substance asulam (variant evaluated asulam-sodium). EFSA J 16, e05251.

Fang, J.P., Zhang, Y.H., Liu, T.T., Yan, B.J., Li, J., and Dong, L.Y. (2019). Target-site and metabolic resistance mechanisms to penoxsulam in barnyardgrass (Echinochloa crus-galli (L.) P. Beauv). J. Agric. Food Chem. 67, 8085–8095.

Fernández-Villa, D., Aguilar, M.R., and Rojo, L. (2019). Folic acid antagonists: antimicrobial and immunomodulating mechanisms and applications. Int. J. Mol. Sci. 20, 4996.

Griffith, E.C., Wallace, M.J., Wu, Y., Kumar, G., Gajewski, S., Jackson, P., Phelps, G.A., Zheng, Z., Rock, C.O., Lee, R.E., and White, S.W. (2018). The structural and functional basis for recurring sulfa drug resistance mutations in Staphylococcus aureus dihydropteroate synthase. Front Microbiol 9, 1369.

Hammoudeh, D.I., Daté, M., Yun, M.K., Zhang, W., Boyd, V.A., Viacava Follis, A., Griffith, E., Lee, R.E., Bashford, D., and White, S.W. (2014). Identification and characterization of an allosteric inhibitory site on dihydropteroate synthase. ACS Chem. Biol. 9, 1294–1302.

Hanson, A.D., and Gregory, J.F. (2002). Synthesis and turnover of folates in plants. Curr. Opin. Plant Biol. 5, 244–249.

Hanson, A.D., and Gregory, J.F. (2011). Folate biosynthesis, turnover, and transport in plants. Annu. Rev. Plant Biol. 62, 105–125.

Heap, I. (2021). The international survey of herbicide resistant weeds.

Hevener, K.E., Yun, M.-K., Qi, J., Kerr, I.D., Babaoglu, K., Hurdle, J.G., Balakrishna, K., White, S.W., and Lee, R.E. (2010). Structural studies of pterin-based inhibitors of dihydropteroate synthase. J. Med. Chem. 53, 166–177.

Hossain, T., Rosenberg, I., Selhub, J., Kishore, G., Beachy, R., and Schubert, K. (2004). Enhancement of folates in plants through metabolic engineering. Proc. Natl. Acad. Sci. USA 101, 5158–5163.

Ji, H., Liu, D., and Yang, Z. (2021). High oil accumulation in tuber of yellow nutsedge compared to purple nutsedge is associated with more abundant expression of genes involved in fatty acid synthesis and triacylglycerol storage. Biotechnol. Biofuels. 14, 1–24.

Krissinel, E., and Henrick, K. (2007). Inference of macromolecular assemblies from crystalline state. J. Mol. Biol. 372, 774–797.

Lawrence, M.C., Iliades, P., Fernley, R.T., Berglez, J., Pilling, P.A., and Macreadie, I.G. (2005). The three-dimensional structure of the bifunctional 6-hydroxymethyl-7,8-dihydropterin pyrophosphokinase/dihydropteroate synthase of Saccharomyces cerevisiae. J. Mol. Biol. 348, 655–670.

Lever, O.W., Bell, L.N., McGuire, H.M., and Ferone, R. (1985). Monocyclic pteridine analogues. Inhibition of Escherichia coli dihydropteroate synthase by 6-amino-5-nitrosoisocytosines. J. Med. Chem. 28, 1870–1874.

Lever, O.W., Bell, L.N., Hyman, C., McGuire, H.M., and Ferone, R. (1986). Inhibitors of dihydropteroate synthase: substituent effects in the side-chain aromatic ring of 6-[[3-(aryloxy)propyl]amino]-5-nitrosoisocytosines and synthesis and inhibitory potency of bridged 5-nitrosoisocytosine-p-aminobenzoic acid analogues. J. Med. Chem. 29, 665–670.

Liu, F., Ying, G.G., Tao, R., Zhao, J.L., Yang, J.F., and Zhao, L.F. (2009). Effects of six selected antibiotics on plant growth and soil microbial and enzymatic activities. Environ. Pollut. 157, 1636–1642.

McIntosh, S.R., Brushett, D., and Henry, R.J. (2008). GTP cyclohydrolase 1 expression and folate accumulation in the developing wheat seed. J. Cereal Sci. 48, 503–512.

Morgan, R.E., Batot, G.O., Dement, J.M., Rao, V.A., Eadsforth, T.C., and Hunter, W.N. (2011). Crystal structures of Burkholderia cenocepacia dihydropteroate synthase in the apo-form and complexed with the product 7,8-dihydropteroate. BMC Struct. Biol. 11, 21.

Mouillon, J.-M., Ravanel, S., Douce, R., and Rébeillé, F. (2002). Folate synthesis in higher-plant mitochondria: coupling between the dihydropterin pyrophosphokinase and the dihydropteroate synthase activities. Biochem. J. 363, 313–319.

Navarrete, O., Van Daele, J., Stove, C., Lambert, W., Van Der Straeten, D., and Storozhenko, S. (2012). A folate independent role for cytosolic HPPK/DHPS upon stress in Arabidopsis thaliana. Phytochemistry 73, 23–33.

Pashley, T.V., Volpe, F., Pudney, M., Hyde, J.E., Sims, P.F., and Delves, C.J. (1997). Isolation and molecular characterization of the bifunctional hydroxymethyldihydropterin pyrophosphokinase-dihydropteroate synthase gene from Toxoplasma gondii. Mol. Biochem. Parasitol. 86, 37–47.

Pemble, C.W., Mehta, P.K., Mehra, S., Li, Z., Nourse, A., Lee, R.E., and White, S.W. (2010). Crystal structure of the 6-hydroxymethyl-7,8-dihydropterin pyrophosphokinase dihydropteroate synthase bifunctional enzyme from Francisella tularensis. PLoS One 5, e14165.

Piotrowicz-Cieślak, A.I., Adomas, B., Nałecz-Jawecki, G., and Michalczyk, D.J. (2010). Phytotoxicity of sulfamethazine soil pollutant to six legume plant species. J. Toxicol. Environ. Health A. 73, 1220–1229.

Pornthanakasem, W., Riangrungroj, P., Chitnumsub, P., Ittarat, W., Kongkasuriyachai, D., Uthaipibull, C., Yuthavong, Y., and Leartsakulpanich, U. (2016). Role of Plasmodium vivax dihydropteroate synthase polymorphisms in sulfa drug resistance. Antimicrob. Agents Chemother. 60, 4453–4463.

Prabhu, V., Lui, H., and King, J. (1997). Arabidopsis dihydropteroate synthase: General properties and inhibition by reaction product and sulfonamides. Phytochemistry 45, 23–27.

Rébeillé, F., Macherel, D., Mouillon, J.-M., Garin, J., and Douce, R. (1997). Folate biosynthesis in higher plants: purification and molecular cloning of a bifunctional 6-hydroxymethyl-7,8-dihydropterin pyrophosphokinase/7,8-dihydropteroate synthase localized in mitochondria. EMBO J. 16, 947–957.

Robert, X., and Gouet, P. (2014). Deciphering key features in protein structures with the new ENDscript server. Nucleic Acids Res. 42, W320–324.

Roland, S., Ferone, R., Harvey, R.J., Styles, V.L., and Morrison, R.W. (1979). The characteristics and significance of sulfonamides as substrates for Escherichia coli dihydropteroate synthase. J. Biol. Chem. 254, 10337–10345.

Schrödinger, L.L.C. (2015). The PyMOL Molecular Graphics System, version 1.8.

Shaw, G.X., Li, Y., Shi, G., Wu, Y., Cherry, S., Needle, D., Zhang, D., Tropea, J.E., Waugh, D.S., Yan, H., and Ji, X. (2014). Structural enzymology and inhibition of the bi-functional folate pathway enzyme HPPK–DHPS from the biowarfare agent Francisella tularensis. FEBS J. 281, 4123–4137.

Shi, G., Shaw, G., Liang, Y.-H., Subburaman, P., Li, Y., Wu, Y., Yan, H., and Ji, X. (2012). Bisubstrate analogue inhibitors of 6-hydroxymethyl-7,8-dihydropterin pyrophosphokinase: New design with improved properties. Bioorg. Med. Chem. 20, 47–57.

Sievers, F., Wilm, A., Dineen, D., Gibson, T.J., Karplus, K., Li, W., Lopez, R., McWilliam, H., Remmert, M., Söding, J., Thompson, J.D., and Higgins, D.G. (2011). Fast, scalable generation of high-quality protein multiple sequence alignments using Clustal Omega. Mol. Syst. Biol. 7, 539.

Storozhenko, S., Navarrete, O., Ravanel, S., De Brouwer, V., Chaerle, P., Zhang, G.-F., Bastien, O., Lambert, W., Rébeillé, F., and Van Der Straeten, D. (2007). Cytosolic hydroxymethyldihydropterin pyrophosphokinase/dihydropteroate synthase from Arabidopsis thaliana: a specific role in early development and stress response. J. Biol. Chem. 282, 10749–10761.

Sukhoverkov, K.V., Corral, M.G., Leroux, J., Haywood, J., Johnen, P., Newton, T., Stubbs, K.A., and Mylne, J.S. (2021). Improved herbicide discovery using physico-chemical rules refined by antimalarial library screening. RSC Adv. 11, 8459–8467.

Tan, K., Kim, Y., Hatzos-Skintges, C., Chang, C., Cuff, M., Chhor, G., Osipiuk, J., Michalska, K., Nocek, B., An, H., Babnigg, G., Bigelow, L., Joachimiak, G., Li, H., Mack, J., Makowska-Grzyska, M., Maltseva, N., Mulligan, R., Tesar, C., Zhou, M., and Joachimiak, A. (2014). Salvage of failed protein targets by reductive alkylation. Methods Mol. Biol. 1140, 189–200.

Terwilliger, T.C., Grosse-Kunstleve, R.W., Afonine, P.V., Moriarty, N.W., Zwart, P.H., Hung, L.-W., Read, R.J., and Adams, P.D. (2008). Iterative model building, structure refinement and density modification with the PHENIX AutoBuild wizard. Acta Crystallogr. D 64, 61–69.

Tickle, I.J., Flensburg, C., Keller, P., Paciorek, W., Sharff, A., Vonrhein, C., Bricogne, G.. (2018). STARANISO (Cambridge, United Kingdom: Global Phasing Ltd.).

Triglia, T., and Cowman, A.F. (1994). Primary structure and expression of the dihydropteroate synthetase gene of Plasmodium falciparum. Proc. Natl. Acad. Sci. USA 91, 7149–7153.

Trott, O., and Olson, A.J. (2010). AutoDock Vina: improving the speed and accuracy of docking with a new scoring function, efficient optimization, and multithreading. J. Comput. Chem. 31, 455–461.

Ulrich, G., Gabriele, J.K., Christoph, H., Adelbert, B., Jörn, K., Dietmar, B., and Johannes, H.H. (2004). Characterization of the Saccharomyces cerevisiae Fol1 protein: starvation for C1 carrier induces pseudohyphal growth. Mol. Biol. Cell 15, 3811–3828.

Waterhouse, A., Bertoni, M., Bienert, S., Studer, G., Tauriello, G., Gumienny, R., Heer, F.T., de Beer, T.A.P., Rempfer, C., Bordoli, L., Lepore, R., and Schwede, T. (2018). SWISS-MODEL: homology modelling of protein structures and complexes. Nucleic Acids Res. 46, W296–w303.

Winn, M. D., Ballard, C.C., Cowtan, K.D., Dodson, E.J., Emsley, P., Evans, P.R., Keegan, R.M., Krissinel, E.B., Leslie, A.G.W., McCoy, A., McNicholas, S.J., Murshudov, G.N. S. Pannu, N., Potterton, E.A., Powell, H.R., Read, R.J., Vagin, A., and Wilson, K.S. (2011). Overview of the CCP4 suite and current developments. Acta Crystallogr. D 67, 235–242.

Yang, J., Yan, R., Roy, A., Xu, D., Poisson, J., and Zhang, Y. (2015a). The I-TASSER Suite: protein structure and function prediction. Nat. Methods 12, 7–8.

Yang, S., Jan, Y.H., Mishin, V., Richardson, J.R., Hossain, M.M., Heindel, N.D., Heck, D.E., Laskin, D.L., and Laskin, J.D. (2015b). Sulfa drugs inhibit sepiapterin reduction and chemical redox cycling by sepiapterin reductase. J. Pharmacol. Exp. Ther. 352, 529–540.

Yogavel, M., Nettleship, J.E., Sharma, A., Harlos, K., Jamwal, A., Chaturvedi, R., Sharma, M., Jain, V., Chhibber-Goel, J., and Sharma, A. (2018). Structure of 6-hydroxymethyl-7,8-dihydropterin pyrophosphokinase-dihydropteroate synthase from Plasmodium vivax sheds light on drug resistance. J. Biol. Chem. 293, 14962–14972.

Yun, M.-K., Hoagland, D., Kumar, G., Waddell, M.B., Rock, C.O., Lee, R.E., and White, S.W. (2014). The identification, analysis and structure-based development of novel inhibitors of 6-hydroxymethyl-7,8-dihydropterin pyrophosphokinase. Bioorg. Med. Chem. 22, 2157–2165.

Yun, M.-K., Wu, Y., Li, Z., Zhao, Y., Waddell, M.B., Ferreira, A.M., Lee, R.E., Bashford, D., and White, S.W. (2012). Catalysis and sulfa drug resistance in dihydropteroate synthase: crystal structures reveal the catalytic mechanism of DHPS and the structural basis of sulfa drug action and resistance. Science 335, 1110–1114.

Zhao, Y., Hammoudeh, D., Yun, M.-K., Qi, J., White, S.W., and Lee, R.E. (2012). Structure-based design of novel pyrimido[4,5-c]pyridazine derivatives as dihydropteroate synthase inhibitors with increased affinity. ChemMedChem. 7, 861–870.

Zhao, Y., Shadrick, W.R., Wallace, M.J., Wu, Y., Griffith, E.C., Qi, J., Yun, M.-K., White, S.W., and Lee, R.E. (2016). Pterin-sulfa conjugates as dihydropteroate synthase inhibitors and antibacterial agents. Bioorg. Med. Chem. Lett. 26, 3950–3954.

